# Phosphatidylserine binding regulates TIM-3 effects on T cell receptor signaling

**DOI:** 10.1101/2021.05.07.443190

**Authors:** Courtney M. Smith, Alice Li, Nithya Krishnamurthy, Mark A. Lemmon

## Abstract

Co-signaling receptors for the T cell receptor are important therapeutic targets, with blocking co-inhibitory receptors such as PD-1 now central in immuno-oncology. Advancing additional therapeutic immune modulation approaches requires understanding ligand regulation of other co-signaling receptors. One poorly understood therapeutic target is TIM-3 (T cell immunoglobulin and mucin domain containing-3). Which ligands are relevant for TIM-3 signaling is unclear, and different studies have reported it as co-inhibitory or co-stimulatory. Here, we show that TIM-3 promotes NF-κB signaling and IL-2 secretion following T cell receptor stimulation in Jurkat cells, and is regulated by phosphatidylserine (PS) binding. TIM-3 signaling is stimulated by PS exposed constitutively in cultured Jurkat cells, and can be blocked by mutating the PS-binding site or by occluding this site with an antibody. We also find that TIM-3 signaling alters CD28 phosphorylation. Our findings help clarify conflicting literature results with TIM-3, and inform its exploitation as a therapeutic target.

## INTRODUCTION

The functional outcome when an antigen engages the T cell receptor (TCR) depends on the activity of a wide range of ‘co-signaling receptors’ in T cells [1], which can be stimulatory (like CD28) or inhibitory (like CTLA-4 and PD-1). The co-stimulatory receptors promote T cell activity and play roles in priming naïve T cells or forming memory T cells. Conversely, co-inhibitory receptors restrain T cell activity and are important for immunological homeostasis – preventing autoimmunity under normal circumstances, but also allowing tumors to evade immune responses in cancer. Both classes of co-receptor have offered important opportunities in immunotherapy, including suppression of co-stimulatory receptor signaling in autoimmunity [2] and suppression of co-inhibitory receptors – or immune checkpoint blockade (ICB) – in cancer [3, 4]. The regulatory ligands are known for most co-signaling receptors currently targeted therapeutically, as is the designation of the receptor as co-inhibitory or co-stimulatory [5]. An exception to this is TIM-3, or T cell immunoglobulin and mucin domain containing-3, for which multiple ligands have been proposed, and both co-stimulatory and co-inhibitory activities have been described [6–8].

TIM-3 was first identified as a marker of CD4 T helper 1 (T_H_1) cells and CD8 cytotoxic T (T_C_1) cells [9], and was later identified on exhausted T cells in chronic viral infection and cancer [10–12]. Early in vivo studies pointed to a co-inhibitory role for TIM-3 [13, 14]. Blocking TIM-3 engagement in mice with antibodies or soluble TIM-3 extracellular domain was found to increase T_H_1 cell proliferation, and TIM-3 deficient mice showed defects in immune tolerance. In vitro studies have variously reached the conclusion that TIM-3 can either suppress or promote T cell signaling [6, 15–19]. Moreover, TIM-3 does not have a definable intracellular ITIM (immunoreceptor tyrosine-based inhibitory motif) or ITSM (immunoreceptor tyrosine-based switch motif) – motifs that normally characterize co-inhibitory receptors and which recruit SH2 domain-containing phosphatases to reduce T cell signaling [20]. Nonetheless, pre-clinical studies have indicated that antibody blockade of TIM-3, in combination with PD-1 blockade, may be a promising therapeutic approach in cancer [21, 22]. Several TIM-3 antibodies are now in clinical trials [23], underlining the need to understand it mechanistically.

Adding further complexity to understanding TIM-3, several regulatory ligands have been reported. The first was the lectin family member galectin-9 [24], which has two β-galactoside-binding carbohydrate-recognition domains. Galectin-9 is thought to induce T cell death by binding to carbohydrates on TIM-3, although some work has refuted this [25, 26]. The glycoprotein CEACAM1/CD66a and the alarmin HMGB1 have also been reported as TIM-3 ligands [8], but their mechanism and relevance are not yet clear. Another major TIM-3 ligand is the membrane phospholipid phosphatidylserine (PS), exposed on the surface of cells undergoing apoptosis and other processes [27, 28], including T cell activation [29]. Homology between TIM-3 and the known PS receptor TIM-4 [30] initially suggested PS as a ligand. Crystallographic and binding studies have confirmed that TIM-3 binds PS, and TIM-3 can also facilitate binding to and engulfment of apoptotic cells (efferocytosis) by macrophages like its relatives TIM-1 and TIM-4 [31–33]. Importantly, however, the role played by PS binding in modulating TIM-3 function in T cells has not been elucidated – although it was recently reported that the epitopes bound by immunomodulatory TIM-3 antibodies all overlap with the PS binding site [34].

Here, we explored the importance of PS for the effect of TIM-3 on TCR signaling, using a Jurkat cell model. We asked whether PS is a key regulatory ligand for TIM-3, beyond the role of this phospholipid in promoting engulfment of apoptotic cells when TIM-3 is expressed on macrophages. In agreement with several previous studies, we observed a co-stimulatory effect when TIM-3 was overexpressed in Jurkat cells. We found that this effect requires the TIM-3 extracellular region, suggesting ligand-dependent regulation. Further, we showed that TIM-3-dependent enhancement of TCR signaling is blocked by mutations that prevent PS binding or by an antibody to the PS-binding site. Thus, endogenous PS in our culture system appears to promote TIM-3 effects on TCR signaling. These findings argue that the co-receptor function of TIM-3 specifically depends on PS binding, which has important mechanistic implications for therapeutic targeting of this receptor.

## MATERIALS AND METHODS

### Cell culture

The NF-κB/Jurkat/GFP^TM^ Transcriptional Reporter cell line was obtained from System Biosciences, and was cultured in RPMI-1640 media supplemented with 10% FBS, 100 U/ml penicillin, and 100 μg/ml streptomycin at 37°C with 5% CO_2_ in a humidified environment. HEK293LTV cells (Cell Biolabs Inc.) used to generate lentivirus were cultured in DMEM supplemented with 10% FBS, 100 U/ml penicillin, and 100 μg/ml streptomycin at 37°C in 5% CO2 in a humidified environment. Human peripheral blood mononuclear cells (PBMCs) were a generous gift from Susan Kaech. PBMCs were cultured in RPMI 1640 media supplemented with 10% FBS, 2 mM GlutaMAX (ThermoFisher Scientific), 50 μM beta-mercaptoethanol, 100 U/ml penicillin, and 100 μg/ml streptomycin at 37°C with 5% CO_2_ in a humidified environment. Raji B cells were obtained from American Type Culture Collection and were cultured in RPMI-1640 media supplemented with 10% FCS, 100 U/ml penicillin, and 100 μg/ml streptomycin at 37°C with 5% CO_2_ in a humidified environment. Expi293F cells used for protein expression (ThermoFisher Scientific) were cultured in Expi293 media (ThermoFisher Scientific) at 37°C in a humidified environment containing 8% CO_2_ while shaking.

### Plasmids

pCDEF3-hTIM-3 was a gift from Lawrence Kane (Addgene plasmid # 49212), and contained the natural variant L119, which was corrected to R119 by site-directed mutagenesis. Full length human TIM-3 (R119) was then sub-cloned into pcDNA3.1(+) using Gibson assembly [35].

Human PD-1 in pENTR223 [36] was a generous gift from Aaron Ring at Yale University. The lentivirus transfer plasmid was a generous gift from the laboratory of Mandar Muzumdar at Yale University and contained AmpR, PuroR under the SV40 promoter, and the gene of interest under the PGK promoter [37]. cDNA fragments encoding human TIM-3 and human PD-1 were sub-cloned into the lentivirus transfer plasmid. The lentivirus envelope and packaging plasmids pMD2.G and pCMV delta R8.2 were gifts from Didier Trono (Addgene plasmids # 12259 and 12263). Plasmids containing human TIM-1 and human TIM-4 were obtained from Origene. For expression of extracellular regions, cDNA fragments encoding the ECRs of human TIM-3, TIM-1, and TIM-4 were sub-cloned into pcDNA3.1(+) vector using Gibson assembly, introducing a C-terminal hexa-histidine tag.

TIM-1-3, TIM-4-3, and PD-1/TIM-3 chimerae were constructed in the lentivirus transfer vector using Gibson assembly. The TIM-3^F40A^ and TIM-3^I96A/M97A^ variants were generated by QuikChange site-directed mutagenesis (Agilent) in pcDNA3.1 (+) for ECR protein expression and in the lentivirus transfer vector for cellular studies. Sequencing was performed to validate all plasmids before use.

### Antibodies

Cells were stained for flow cytometry analysis with phycoerythrin-conjugated forms of anti-TIM-3 (R & D Systems, catalog #FAB2365P), anti-PD-1 (BioLegend, catalog #329905), anti-TIM-1 (BioLegend, catalog #353903), anti-TIM-4 (BioLegend, catalog #354003), anti-PD-L1 (also labeled with Dazzle 594: BioLegend, catalog #329731), or with an unlabeled TIM-3 primary antibody (R & D Systems, catalog #AF2365) using a phycoerythrin-conjugated anti-goat secondary antibody for detection (R & D Systems, catalog # F0107). PBMCs were stained with phycoerythrin-Dazzle694-labeled anti-CD3 (Biolegend, catalog #317345), PerCP/Cy5.5^TM^-labeled anti-CD8 (Biolegend, catalog #301031), BB515-labeled anti-CD4 (BD Biosciences, catalog #564420), and phycoerythrin-labeled anti-TIM-3 (R & D Systems, catalog #FAB2365P), Western blot analysis was performed with anti-Zap70 pY319 (Cell Signaling Technology, catalog #2701), anti-LAT pY191 (Cell Signaling Technology, catalog #3584), anti-PLCγ pY783 (Cell Signaling Technology, catalog #14008), anti-ERK1/2 pT202/pY204 (Cell Signaling Technology, catalog #9106), anti-Grb2 (Cell Signaling Technology, catalog #3972), anti-CD28 pY218 (Sigma Aldrich, catalog #SAB4504133), anti-CD28 pY191 (Cell Signaling Technology, catalog #16399), anti-AKT pT308 (Cell Signaling Technology cat# 2965), anti-TIM-3 (R & D Systems, catalog #AF2365), anti-TIM-3 (Abcam, catalog #ab241332), and anti-PD-1 (Cell Signaling Technology, catalog #86163). For TCR stimulation, anti-CD3 clone OKT3 (BioLegend, catalog #317326) and anti-CD28 clone CD28.2 (BioLegend, catalog #302943) were used. For functional studies, anti-TIM-3 clone F38.2E2 (ThermoFisher Scientific, catalog #16-3109-85) was used.

### Lentivirus transduction

Lentiviruses were generated by co-transfecting HEK293 LTV cells with lentivirus transfer, envelope, and packaging plasmids using the Mirus Bio TransIT-Lenti Transfection Reagent, according to the manufacturer’s protocol. Briefly, HEK293 LTV cells were cultured to 80-95% confluency. Plasmids were mixed with Opti-MEM and TransIT-Lenti Transfection Reagent and were incubated at room temperature for 10 min before being added to HEK293 LTV cells and incubating at 37°C, 5% CO_2_. After 48 h, virus-containing medium was collected and filtered through a 0.22 μm filter. Filtered virus suspension was then added to Jurkat cells (2 x 10^5^ cells), and cells were transduced with lentivirus by spinfection (spinning at 800 x g for 30 min at 32^°^C) ‘spinoculation’ [38] in sealed aerosol containment lids. Three days after transfection, medium was replaced and cells were selected with puromycin (1 μg/ml) for 2 days. After selection, medium was replaced again and cells were allowed to recover. Following selection, receptor expression was verified by flow cytometry. Cells were washed in FACS Buffer: 10% FBS plus 0.1% NaN_3_ in phosphate-buffered saline (PBS) 0.22 μm filtered. They were then stained with antibodies in the dark. Cells were rewashed and then analyzed on a FACSMelody (Becton Dickinson), with a four-color set up (488 nm and 561 nm lasers with 527/32, 700/54, 582/15, and 613/18 filter sets).

### Expression of TIM-3 on peripheral blood mononuclear cells (PBMCs)

After overnight recovery, thawed PBMCs were transferred to a 24-well plate coated with 1 μg/ml αCD3, and soluble αCD28 (1 μg/ml) was added to the culture to promote outgrowth of T cells. Cells were maintained between 0.25 and 0.5 x 10^6^ cells per cm^2^ with αCD3/αCD28 stimulation for 7 days to promote expression of TIM-3 [39]. After 7 days, PBMCs were washed in FACS Buffer (10% FBS plus 0.1% NaN_3_ in phosphate-buffered saline (PBS) 0.22 μm filtered). PBMCs were then stained with labeled anti-CD3, anti-CD8, anti-CD4, and anti-TIM-3. Cells were then washed again in FACS Buffer before analysis on a FACSMelody. PBMCs were first subjected to doublet discrimination before analyzing for expression of cell surface markers, as shown in Supplementary Figure 1.

### NF-**κ**B reporter assay

NF-κB GFP reporter Jurkat cells were resuspended in RPMI-1640 media with 0% FBS and were serum starved for 4 h. After starvation, cells were counted and 5 x 10^5^ cells were plated in a V-bottom 96-well plate and stimulated with αCD3/αCD28 (1 μg/ml each, or 0.5 μg/ml for αCD28), with additional treatments as stated in figure legends. In experiments investigating effects of anti-TIM-3 (F38.2E2), cells were pre-treated with TIM-3 antibody for 1 h before stimulation with αCD3/αCD28. Cells were returned to the incubator at 37°C, 5% CO_2_ for 16 h. Cells were then washed in FACS Buffer, stained with antibodies, and rewashed prior to analysis using the FACSMelody. Data were analyzed using FlowJo software. GFP signal was quantified for single cells, as determined through SSC-H vs. SSC-W and FSC-H vs. FSC-W gates.

### Analysis of IL-2 secretion

NF-κB GFP reporter Jurkat cells were resuspended in RPMI-1640 media with 0% FBS and were serum starved for 4 h. After starvation, cells were counted and 5 x 10^5^ cells were plated in a V-bottom 96-well plate. Cells were stimulated with 1 μg/ml αCD3 (BioLegend, catalog #317326) plus 0.5 or 1 μg/ml αCD28 (BioLegend, catalog #302943), and returned to the incubator at 37°C, 5% CO_2_, for 16 h. In experiments analyzing the effect of anti-TIM-3 F38.2E2, cells were pre-treated with anti-TIM-3 for 1 h before stimulation with αCD3/αCD28. In experiments using Raji B cell-based stimulation, Raji B cells were loaded with 30 ng/ml Staphylococcal enterotoxin E (Toxin Technology Inc., catalog #ET404) for 30 min at 37°C and were washed in serum-free RPMI-1640 media to remove excess toxin. Jurkat NF-κB GFP reporter cells were serum starved in RPMI-1640 media with 0% FBS for 3 h. After starvation, 2 x 10^5^ Jurkat cells were mixed with 10^5^ SEE-loaded Raji B cells per well in a 96-well U-bottom plate, as described [40]. The plate was centrifuged for 1 min at 300 x g to initiate cell contact, and plate was returned to incubator at 37°C, 5% CO_2_, for 6 h. Following antibody or cell-based stimulation, media was collected for analysis of IL-2 secretion by ELISA (R & D Systems, cat #D2050), following the IL-2 Quantikine ELISA manual. Experiments were performed in technical duplicate and biological triplicate.

### Western blot analysis

NF-κB GFP reporter Jurkat cells were serum starved and stimulated as above, with additional treatments as stated in figure legends. Where wortmannin was used (in Figure 2C), cells were pre-treated with drug for 1 h before stimulation. Cells were pelleted by centrifugation (800 x g for 1.5 min) in the last 1.5 minutes of stimulation. Supernatant was removed, and cells were lysed in ice cold Lysis Buffer (Cell Signaling Technology) with freshly added cOmplete protease inhibitor (Roche) and PhosSTOP phosphatase inhibitor (Roche). Lysates were clarified by centrifugation (10,000 x g for 10 min at 4°C), and clarified lysate was isolated. Lysates were mixed with NuPAGE LDS Sample Buffer (Invitrogen) plus dithiothreitol (DTT) and were boiled for 8 min. Samples were then analyzed by SDS-PAGE using NuPAGE 4-12% Bis-Tris gels. Gels were transferred to 0.22 μm nitrocellulose membranes using the Xcell Surelock Electrophoresis system (ThermoFisher Scientific). To probe multiple proteins simultaneously without requiring stripping and reprobing, the multistrip Western blotting procedure was used [41], in which horizontal strips are excised from the nitrocellulose membrane corresponding to molecular weight guides to allow probing of one blot with several antibodies. Membranes were blocked with 4% BSA in Tris-buffered saline plus Tween20 (TBS-T), probed with primary antibody for 1 h to overnight, and then probed with HRP-tagged secondary antibodies for 1 h. Detection was by enhanced chemiluminescence using SuperSignal Western Pico PLUS Chemiluminescent substrate (ThermoFisher Scientific), with visualization and quantitation using a Kodak Image Station (Kodak Scientific).

### Detection of Annexin V staining by flow cytometry

PerCP-Cy^TM^5.5 or FITC-labeled Annexin V were obtained from BioLegend (catalog #640936 and 640905). As specified by the manufacturer, cells were washed twice with ice cold PBS and then resuspended in 10 mM HEPES pH 7.5, 140 mM NaCl, 2.5 mM CaCl_2_ (binding buffer).

Annexin V-PerCP-Cy5.5 or -FITC was added to each sample and incubated for 15 min in the dark. Additional binding buffer was added to each sample, followed by analysis using FACSMelody (Becton Dickinson) and FlowJo software.

### Detection of Annexin V staining by widefield fluorescence microscopy

AlexaFluor® 647-labeled Annexin V was obtained from Biolegend (catalog #640911). As detailed above, Jurkat NF-κB GFP reporter cells were washed twice with ice cold PBS and resuspended in Annexin V binding buffer (10 mM HEPES pH 7.5, 140 mM NaCl, 2.5 mM CaCl_2_). Annexin V-AlexaFluor® 647 was added to cells and incubated for 15 minutes in the dark. Cells were then fixed in 4% paraformaldehyde in PBS with 2.5 mM CaCl_2_ for 15 minutes. Annexin V-stained cells were cytospun onto microscope slides at 800 RPM for 5 minutes. Cells were fixed with ProLong^TM^ Diamond Antifade Mountant with DAPI (ThermoFisher Scientific, catalog #P36966) per the manufacturer’s instructions. Images were collected with a Nikon Eclipse Ti2 widefield fluorescence microscope using a Nikon Plan Apo λ 40x 0.95 N.A. objective. Excitation was provided by a Sola light engine with a DAPI or Cy5 filter cube sets.

Emission was collected by an sCMOS pco.edge camera. Images were analyzed with FIJI software.

### Protein expression and purification

As specified in the Expi293 Expression System manual, Expi293F cells were cultured to reach a density of 4.5-5.5 x 10^6^ cells/ml on the day of transfection, and seeded at 3 x 10^6^ cells/ml. A transfection mixture containing DNA vector (1 μg per ml of culture to be infected), Expifectamine reagent (ThermoFisher Scientific), and Opti-MEM was then incubated at room temperature for 10 min and subsequently added to the culture. Cells were returned to the humidified incubator at 37°C, 8% CO_2_. Expi293 enhancers (ThermoFisher Scientific) were added 18-22 h after transfection, and the culture was harvested 4-6 days later by centrifugation (1,000 RPM). Culture supernatant was collected and diafiltered against 4 times the culture volume with 10 mM HEPES pH 8, 150 mM NaCl (TIM Buffer 1). Nickel affinity chromatography was performed on the diafiltered supernatant with Ni-NTA agarose (Qiagen), eluting fractions with increasing concentrations of imidazole in TIM Buffer 1. Protein-containing fractions were pooled, diluted to 50 mM NaCl, filtered (0.22 μm), and injected onto a Fractogel TMAE column (EMD Millipore) equilibrated in 25 mM HEPES pH 8 at 50 mM NaCl. Protein was eluted using a gradient from 50 to 700 mM NaCl in 25 mM HEPES pH 8. Peak fractions (∼150 mM NaCl) were collected, concentrated using a 10 kDa MWCO centrifugal filter (EMD Millipore), filtered (0.22 μm), and applied to a Superose 12 10/300 column (Cytiva Life Sciences) for gel filtration in TIM Buffer 2 (10 mM HEPES pH 7.6, 150 mM NaCl). Peak fractions were collected, concentrated using a 10 kDa MWCO centrifugal filter (EMD Millipore) and filtered (0.22 μm). SDS-PAGE was performed to confirm protein size and purity. Protein concentration was determined by 280 nm absorbance using a NanoDrop spectrophotometer (ThermoFisher Scientific) with calculated extinction coefficients.

### Vesicle preparation

Lipids were purchased from Avanti Polar Lipids in chloroform solution, including dioleoylphosphatidylcholine (DOPC), dioleoylphosphatidylserine (DOPS), dioleoylphosphatidic acid (DOPA), and dioleoylphosphatidylethanolamine (DOPE). Lipid solutions were combined at the appropriate molar ratios in a glass vial. Chloroform was blown off under a stream of nitrogen gas, fully drying the lipid mixture under vacuum. Lipid mixtures were rehydrated with 10 mM HEPES pH 7.6, 150 mM NaCl, vortexed to mix, and subjected to at least 10 freeze-thaw cycles, where the suspension was frozen in liquid nitrogen and thawed in a warm sonicating water bath, to generate unilamellar vesicles. Vesicle suspensions were stored at −20°C. Before use, thawed sonicated vesicles were extruded using a 100 nm filter membrane in an Avanti Mini Extruder.

### Surface plasmon resonance studies

Surface plasmon resonance (SPR) analyses of protein-lipid interactions were performed using a Biacore 3000 instrument as described [42, 43]. Lipid vesicles containing DOPC or the specified percentage (mole/mole) of lipid in a DOPC background were immobilized by flowing lipid vesicles across an L1 chip (Cytiva) in TIM Buffer 2 (10 mM HEPES pH 7.6, 150 mM NaCl). After immobilization, purified proteins were flowed across the chip at varying concentrations in the presence or absence of 1 mM CaCl_2_. Resonance units detected by the Biacore 3000 were corrected for background (DOPC) binding on a separate sensorchip surface, and were plotted against protein concentration. Curves were fit using the equation: RU_max_ = B_max_ x [TIM]/ (K_d_ + [TIM]), and apparent K_d_ was estimated from these curves.

### Quantification and statistical analysis

Experiments were completed with at least three biological repeats unless noted otherwise. For transcriptional reporter experiments, each independent experiment included three technical repeats. In addition to mean fluorescence intensity, the percent of GFP-positive cells and the median fluorescence intensity of GFP-positive cells were quantitated for transcriptional reporter experiments, to ascertain whether analysis method influenced interpretation. In all cases, the result was the same as seen when assessing mean fluorescence intensity as done in the figures – lending confidence to our analysis and its qualitative conclusions. All quantitated values are represented as mean values ± standard deviation. Statistical analysis was performed with two-tailed, unpaired Student’s t-test.

## RESULTS

### TIM-3 promotes TCR signaling when overexpressed in Jurkat cells

In considering possible mechanisms of TIM-3 signaling, a key question to resolve [8] is whether TIM-3 functions in T cells as a co-inhibitory or co-stimulatory receptor. In vitro studies have suggested both. Lee et al. (2012) reported that elevating TIM-3 expression in Jurkat cells inhibits the NFAT pathway. Moreover, Tomkowicz et al. (2015) reported that TIM-3 overexpression suppressed NFAT and NF-κB signaling following TCR activation with αCD3/αCD28. In stark contrast, Lee et al. (2011) reported that expressing either murine TIM-3 or human TIM-3 in Jurkat cells had the opposite effect – promoting both NF-κB and NFAT signaling. Other studies have also supported a positive role for TIM-3 in acute activation of murine CD8 T cells [6, 16]. Thus, these and other studies argue that the influence of TIM-3 can be positive or negative depending on its T cell type and possibly its expression level [44]. Moreover, Avery et al. (2018) posited that, in some settings, TIM-3 is more similar to co-stimulatory receptors than to inhibitory receptors like PD-1.

To establish a system for studying TIM-3 regulation, we asked how its overexpression affects TCR activation in Jurkat cells – a useful model system for studying TCR signaling [45]. We first established that TIM-3 was undetectable by flow cytometry or Western blotting in Jurkat cells (Figure 1A) – in agreement with others [17] – and was not inducible by TCR stimulation. We then used lentiviral delivery of TIM-3 to generate stably expressing (TIM-3^+^) derivatives of an NF-κB/Jurkat/GFP^TM^ transcriptional reporter cell line in which NF-κB activation downstream of TCR stimulation promotes GFP expression. Importantly, we confirmed that levels of TIM-3 seen following viral transduction of Jurkat NF-κB GFP Reporter cells were similar to that observed in human primary T cells (Supplementary Figure 1).

**FIGURE 1.**
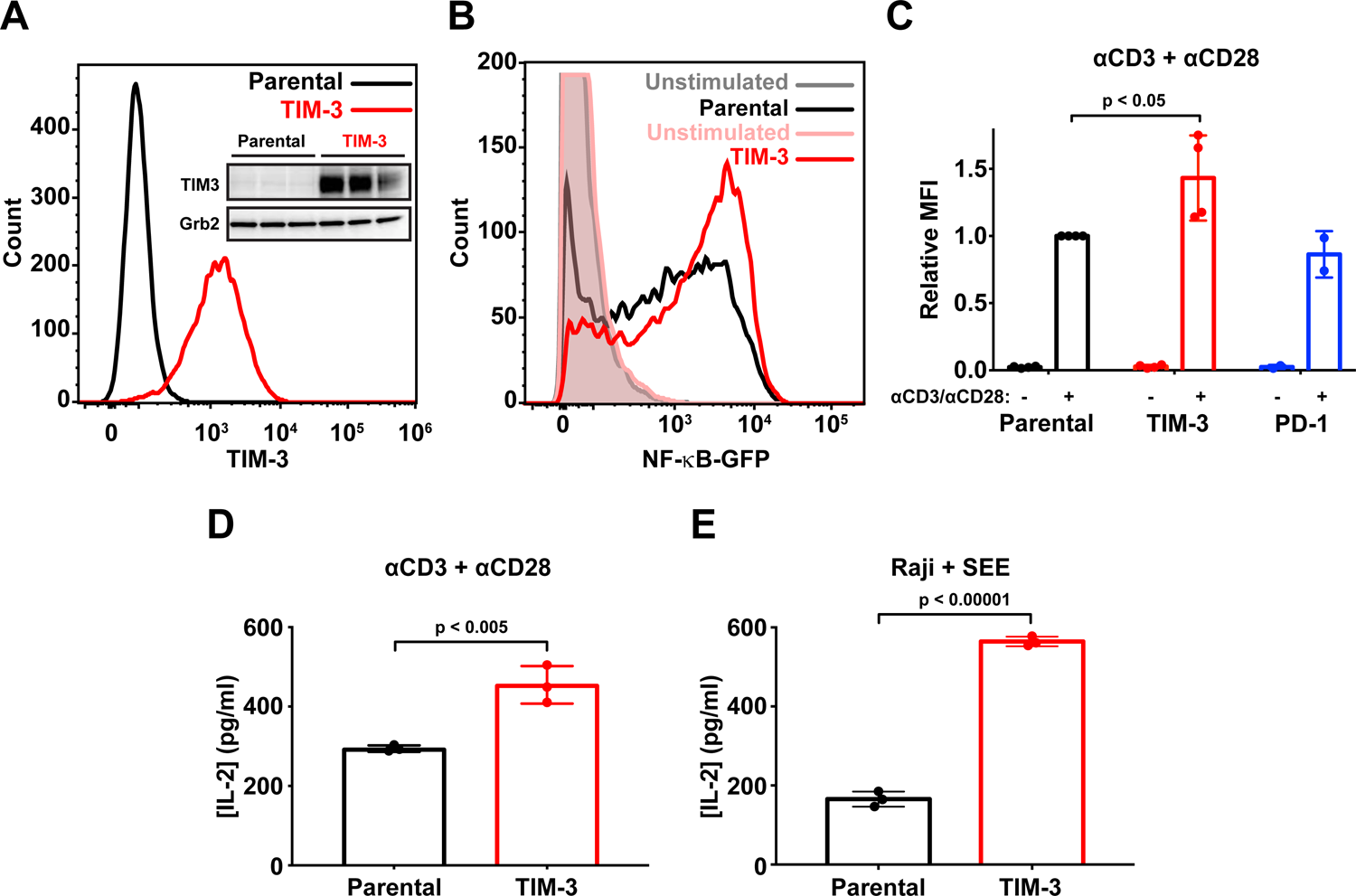
Ectopic expression of TIM-3 promotes activation of Jurkat cells following stimulation. (A) Flow cytometry analysis of human TIM-3 expression in parental (black) and TIM-3 lentivirus-transduced NF-κB GFP reporter Jurkat cells (red), detected with phycoerythrin-conjugated TIM-3 antibody. The inset shows Western blot analysis of TIM-3 expression in the same cells as marked – with three biological repeats of each. (B) A representative histogram of NF-κB-driven GFP expression in stimulated NF-κB reporter cells (black), and those expressing TIM-3 (red), compared with unstimulated parental (grey) and TIM-3-expressing (pink) cells. Stimulation employed 1 μg/ml αCD3 plus 1 μg/ml αCD28 for 16 h to mimic antigen binding, and T cell activation was quantified by flow cytometry analysis of the GFP reporter of NFκB transcriptional activity. Enhanced T cell activation in TIM-3-expressing cells requires cell stimulation, and no change is observed in unstimulated cells. (C) Mean GFP fluorescence intensity (MFI) values for NF-κB GFP reporter assays in cells stimulated as in (**B**) relative to parental NF-κB reporter cells (black). Data represent averages of 4 biological repeats for TIM-3-expressing cells (red) and two for PD-1-expressing cells (blue). The p value for the TIM-3/parental comparison was determined with a two-tailed, unpaired Student’s t-test. (D) ELISA analysis of IL-2 secretion into the medium of parental and TIM-3^+^ NF-κB reporter cells following stimulation as in (**B**). Data represent the mean ± standard deviation (SD) across 3 biological repeats, with p value determined using a two-tailed, unpaired Student’s t-test. (E) Production of IL-2 from parental and TIM-3-expressing Jurkat NF-κB/GFP reporter cells was quantified by ELISA following stimulation by SEE-loaded Raji B cells to mimic formation of the immunological synapse during T cell activation. Data represent mean ± standard deviation (SD) for 3 biological repeats, with p value determined using a two-tailed, unpaired Student’s t-test.

Stimulating NF-κB reporter Jurkat cells with antibodies against CD3 and CD28 (αCD3/αCD28) to mimic antigen binding and TCR activation promoted NF-κB-dependent GFP expression as expected [46]. The GFP signal was significantly increased with TIM-3 expression only after T cell stimulation (Figures 1B,C), consistent with key previous studies summarized above [17] and suggesting a co-stimulatory role for TIM-3 in Jurkat cell activation. By contrast, PD-1 overexpression in the same NF-κB/Jurkat/GFP^TM^ cells had no significant effect on TCR-activated NF-κB signaling (Figure 1C), as expected in the absence of PD-L1 to engage PD-1 in these experiments (Supplementary Figure 2A). In parallel experiments, also consistent with previous studies [17], ELISA assays showed that TIM-3 expression substantially increased the level of IL-2 production induced by TCR activation with αCD3/αCD28 (Figure 1D) – again supporting a co-stimulatory role for TIM-3.

We also stimulated Jurkat cells with Staphylococcal enterotoxin E (SEE)-loaded Raji B cells as antigen-presenting cells to engage the TCR and other key signaling molecules in an immunological synapse (IS). TIM-3 was recently reported to be recruited to the IS during Raji B cell activation of Jurkat T cells [47]. As we observed with antibody stimulation alone, Raji cell activation of Jurkat cells expressing TIM-3 caused them to secrete elevated levels of IL-2 compared to parental cells (Figure 1E), showing that the ability of TIM-3 to promote T cell activation is preserved with IS formation. Together, these data support a co-stimulatory role of TIM-3 in Jurkat T cells when activated by antibodies or by cell-based methods.

### Effects of TIM-3 overexpression on signaling responses to **α**CD3/**α**CD28

To investigate how TIM-3 expression might modulate T cell signaling, we used Western blotting to monitor phosphorylation of components associated with TCR activation after stimulation with αCD3/αCD28. Few differences could be detected when assessing phosphorylation of Zap70, PLCγ1, ERK1/2, or LAT in parental and TIM-3^+^ cells (Supplementary Figure 3). Some experiments did suggest that phosphorylation of Zap70 and LAT is slightly greater and more sustained in TIM-3 overexpressing cells (Supplementary Figures 3B,E), but this did not reach statistical significance. Levels and dynamics of phosphorylation of these proteins also appeared essentially unchanged by PD-1 overexpression (Supplementary Figure 3).

One difference that did reach significance (Figure 2) was seen in the level of CD28 phosphorylation, which was modestly increased in TIM-3 overexpressing cells. This effect was seen for both Y191 phosphorylation (Figure 2A) and Y218 (Figure 2B) at the 5-min time point. Elevated phosphorylation was seen at both sites following αCD3/αCD28 stimulation in parental cells, as expected [48], but TIM-3 expression increased the magnitude of the response by ∼50% – with no apparent elevation of baseline CD28 phosphorylation. Intriguingly, Y191 phosphorylation also seemed to be significantly longer-lived in TIM-3^+^ cells (Figure 2A, Supplementary Figure 3F). Phosphorylation of Y218 in CD28 has been reported to be important for NF-κB activation by CD3/CD28 ligation in Jurkat and primary CD4 T cells [49], consistent with our results for NF-κB activation. Moreover, phosphorylation of all three distal tyrosines in the CD28 tail (including Y218) appears to be required for stimulation of IL-2 secretion by T cells [50]. Thus, alterations in the extent of CD28 phosphorylation and/or its dynamics may be important for the increased NF-κB signaling and IL-2 secretion seen when TIM-3 is overexpressed in Jurkat cells. Phosphorylation of Y191 in CD28 promotes recruitment of phosphoinositide 3-kinase (PI3K) to CD28 [51, 52], and the consequences of altering its phosphorylation are likely blunted in Jurkat cells, which lack PTEN [53]. We did consider that the constitutively high levels of PI3K products seen in Jurkat cells might be responsible for the change in CD28 phosphorylation. However, as shown in Figure 2C, the effect of TIM-3 expression on levels of CD28 phosphorylation was maintained in the presence of the PI3K inhibitor wortmannin – indicating no dependence on PI3K activity.

**FIGURE 2.**
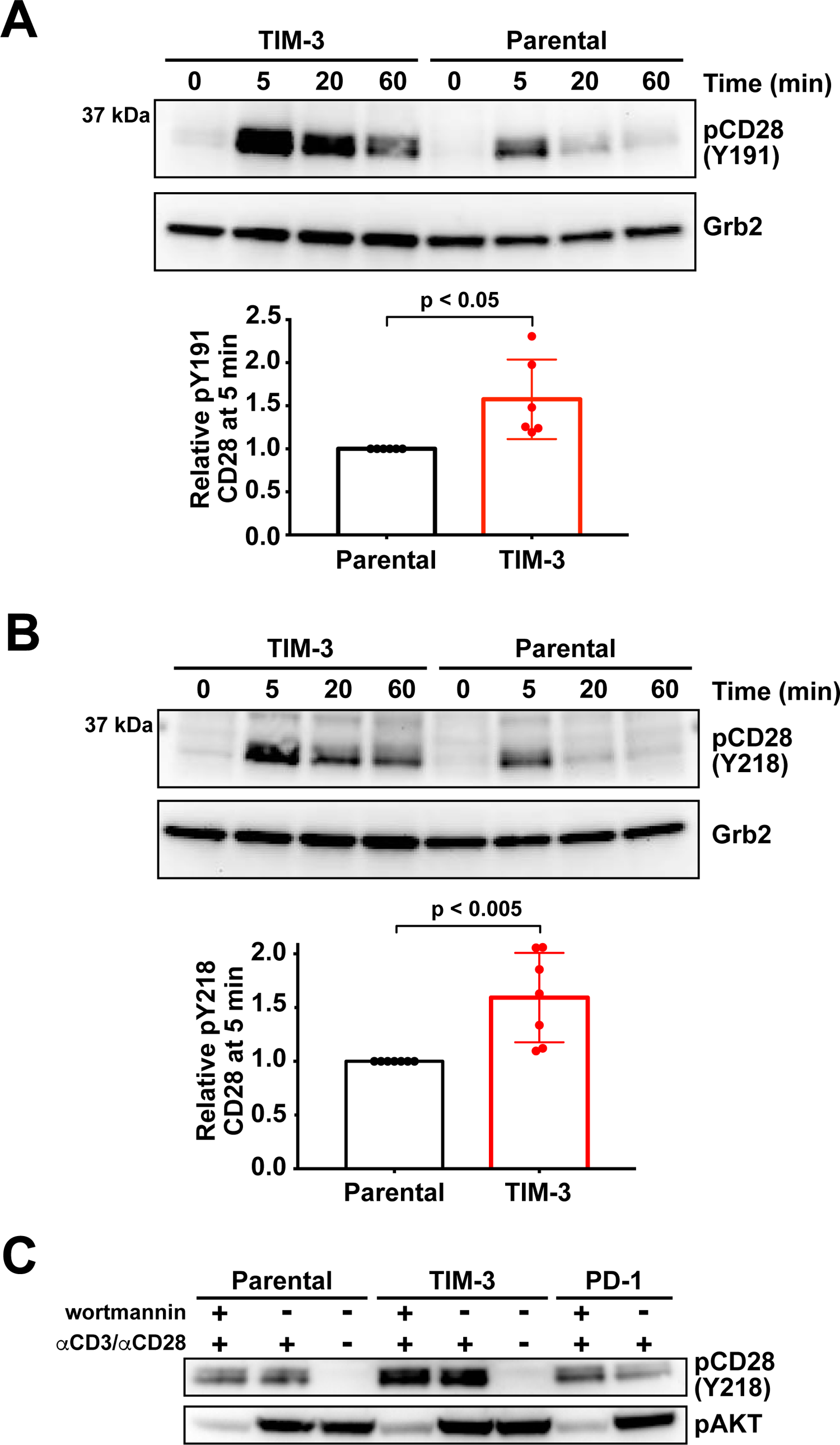
TIM-3 modulates phosphorylation of CD28 after. α**CD3/**α**CD28 stimulation** Western blotting was used to monitor changes in phosphorylation of CD28 following TCR activation with αCD3/αCD28 as in Fig. 1 for 0, 5, 20, or 60 min. (A) Representative Western blot of phospho-CD28 at pY191 for TIM-3-expressing and parental NF-κB reporter Jurkat cells demonstrating elevated phosphorylation of Y191 in TIM-3-expressing cells. Grb2 was used as a loading control as described [41]. Quantification bar graphs indicate mean band intensity ± standard deviation (n=6 biological repeats). Two-tailed, unpaired Student’s t-test was used to determine p values. (B) Representative Western blot of CD28 phosphorylation at Y218 for TIM-3^+^ cells and parental NF-κB reporter cells showing elevated phosphorylation at Y218 at 5 min in TIM-3^+^ cells, with quantitation below the blot. Grb2 was used as a loading control, as above. Bars indicate mean band intensity ± standard deviation for the 5 min time point (7 biological repeats) with p values determined by two-tailed, unpaired Student’s t-test. (C) Parental, TIM-3-expressing, and PD-1-expressing NF-κB reporter Jurkat cells were stimulated for 5 min with αCD3/αCD28 (1 μg/ml each) with- or without the PI3K inhibitor, wortmannin (0.2 μM) to show that PTEN deficiency does not account for the effect of TIM-3 expression on CD28 phosphorylation.

### The TIM-3 extracellular region is required for its activity in Jurkat cells

We next chose to exploit the ability of TIM-3 to enhance NF-κB signaling in the GFP-reporter Jurkat cell line to investigate how the extracellular region of TIM-3 regulates its function, and to understand which of TIM-3’s ligand(s) are important. Lee et al. (2011) previously used a similar strategy to investigate signaling requirements of the TIM-3 cytoplasmic tail. They reported the surprising finding that deletion of the TIM-3 extracellular region did not abolish its ability to enhance αCD3/αCD28-induced NFAT activation [17]. As summarized below, our results contrast with this finding and suggest an important regulatory function for the extracellular region.

Since NF-κB signaling in the Jurkat reporter cell line was increased by expression of TIM-3, but not of PD-1, we asked whether PD-1/TIM-3 chimerae would preserve this effect – which they should if the TIM-3 intracellular region is sufficient for signaling. We replaced the TIM-3 extracellular region with that from PD-1, in ‘PTT’ or ‘PPT’ chimerae (Figure 3A) that retain either the TIM-3 (PTT) or PD-1 (PPT) transmembrane (TM) domain. After generating Jurkat reporter cells stably expressing these chimerae at levels similar to full-length TIM-3 (see Supplementary Figures 2B,E), we assessed their ability to enhance NF-κB activation by αCD3/αCD28. As shown in Figures 3B and C, PTT and PPT cells displayed a slight increase in NF-κB activation compared to parental and PD-1 reporter cells, but not to the same extent as TIM-3 cells, arguing that the TIM-3 extracellular region does play an important role in its signaling effects in Jurkat cells. One possibility is that TIM-3 interacts constitutively with the TCR or other cellular components to influence the effects of TCR stimulation. Alternatively, it is possible – and perhaps more likely – that TIM-3 is engaged by one of its extracellular ligands in Jurkat cell cultures. Indeed, it was previously shown that a biotinylated form of the soluble TIM-3 extracellular region associates with the surface of CD4 T cells in flow cytometry studies [13, 14], and we know that the TIM-3 ligand PS is exposed on the surface of activated T cells, following antigen recognition [29], consistent with this possibility. Interestingly, expression of PTT promoted NF-κB signaling to a slightly greater extent than PPT, possibly implicating the TIM-3 TM domain as recently suggested by Kane and colleagues [47].

**FIGURE 3.**
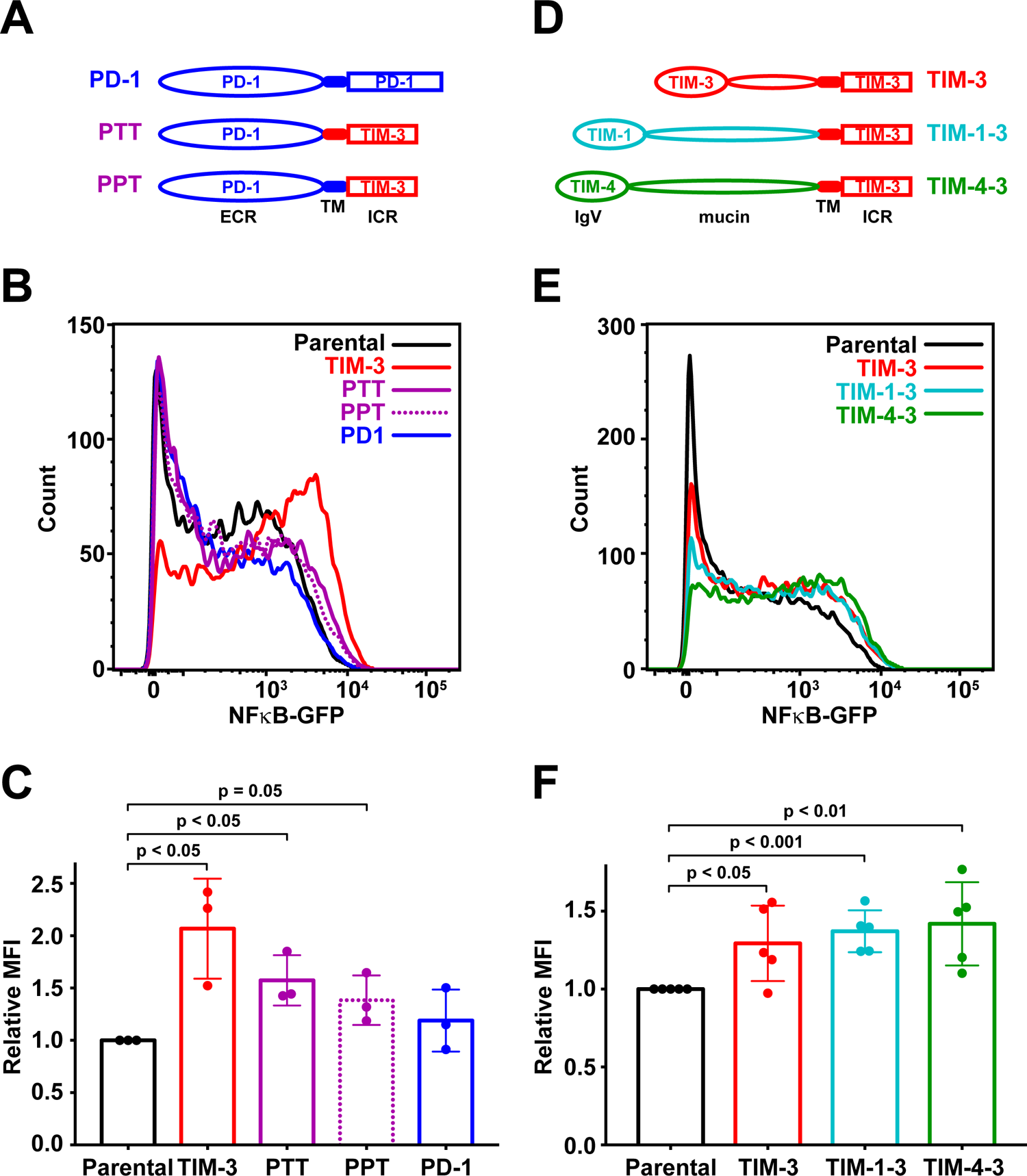
Chimerae with PS-binding extracellular regions promote T cell signaling. (A) Schematic depicting PD-1/TIM-3 chimeric constructs. PTT contains the PD-1 extracellular region (ECR) plus transmembrane (TM) and intracellular regions of TIM-3. In PPT, only the intracellular region is replaced with that from TIM-3. (B) A GFP reporter was used to measure NF-κB transcriptional activity downstream of TCR activation in parental NF-κB reporter Jurkat cells (black) and the same cells engineered to stably express TIM-3 (red curve), PTT (purple solid curve), PPT (purple dotted curve), or PD-1 (blue curve), stimulated with 1 μg/ml αCD3 plus 1 μg/ml αCD28. Representative histograms of NF-κB-driven GFP expression are shown. (C) The relative mean GFP fluorescence intensity (MFI) from 3 biological repeats of the experiment shown in (**B**) is plotted, expressed as fold change above that measured for parental NF-κB GFP reporter cells. Bars are color coded as in (**B**) and represent mean MFI value (± SD) – with p values determined by two-tailed, unpaired Student’s t-test. (D) Schematic depicting TIM-1/TIM-3 and TIM-4/TIM-3 chimeric constructs in which the TIM-3 ECR has been replaced with that of TIM-1 (TIM-1-3) or TIM-4 (TIM-4-3). (E) Representative histogram of NF-κB-driven GFP expression in parental NF-κB GFP reporter cells (black), and those expressing TIM-3 (red), TIM-1-3 (teal), or TIM-4-3 (green) following TCR stimulation with 1 μg/ml αCD3 plus 0.5 μg/ml αCD28 for 16 h. (F) Relative mean GFP fluorescence intensity (MFI) values, as fold change above parental cells in each experiment, is plotted for 5 biological repeats of the experiment shown in (**E**). Bars represent mean ± SD, with p values determined by two-tailed, unpaired Student’s t-test.

### Extracellular regions of other TIM family members can substitute for that of TIM-3

As a first step in understanding the importance of engagement of the TIM-3 extracellular region by its ligands (and what it/they might be), we generated a second set of chimeric receptors that should retain affinity for phosphatidylserine (PS) but not for other proposed TIM-3 ligands. We replaced the TIM-3 extracellular region, which contains the PS-binding pocket, with that of TIM-1 or TIM-4 (which also bind PS – see below), generating the TIM-1-3 and TIM-4-3 chimerae (Figure 3D). These chimerae retain the TIM-3 transmembrane domain and intracellular region. They should not bind to galectin-9, CEACAM1, or HMGB1 [8, 54], and would be unlikely to retain specific TIM-3 extracellular interactions with TCR components. After generating Jurkat NF-κB reporter cells stably expressing these chimeric receptors on the cell surface (Supplementary Figures 2C-E), we asked if they retained the ability to increase αCD3/αCD28-stimulated NF-κB activation. As shown in Figures 3E and F, TIM-1-3 and TIM-4-3 enhanced NF-κB activation to a similar extent as seen for TIM-3 in parallel experiments.

Chimerae with the TIM-3 intracellular region that retain extracellular PS-binding thus seem to preserve the ability to enhance TCR signaling seen for TIM-3.

### Phosphatidylserine exposed in Jurkat cell culture may engage TIM-3

We next asked if TIM-3 signaling could be modulated by adding its putative ligand, PS. We began by quantifying αCD3/αCD28-stimulated NF-κB activity in TIM-3^+^ cells after adding unilamellar vesicles containing high levels of PS. Vesicles containing 20% (mole/mole) dioleoylphosphatidylserine (DOPS) in a background of dioleoylphosphatidylcholine (DOPC) had no effect on NF-κB activity in parental Jurkat cells, but also failed to modulate NF-κB activity in cells expressing TIM-3 to a statistically significant degree when compared with pure PC vesicles (Supplementary Figures 4A,B). We therefore hypothesized that Jurkat cells in culture might expose PS on their surface, which could correspond to the previously observed TIM-3 ligand expressed on CD4 T cells [13, 14]. Indeed, *E. coli*-derived recombinant murine TIM-3 IgV domain was shown to bind T cells, fibroblasts and macrophages from several species [55] – consistent with recognition of a conserved component of these cells, such as PS. As a specific test of the hypothesis that PS on the surface of Jurkat cells promotes TIM-3 effects in our studies, we asked whether adding excess annexin V – a known PS-binding protein – might block surface-exposed PS and suppress TIM-3-dependent elevation of NF-κB activity. Unfortunately, adding annexin V elevated TCR-driven NF-κB signaling independently of TIM-3 expression (Supplementary Figures 4C,D), making this experiment uninterpretable.

**FIGURE 4.**
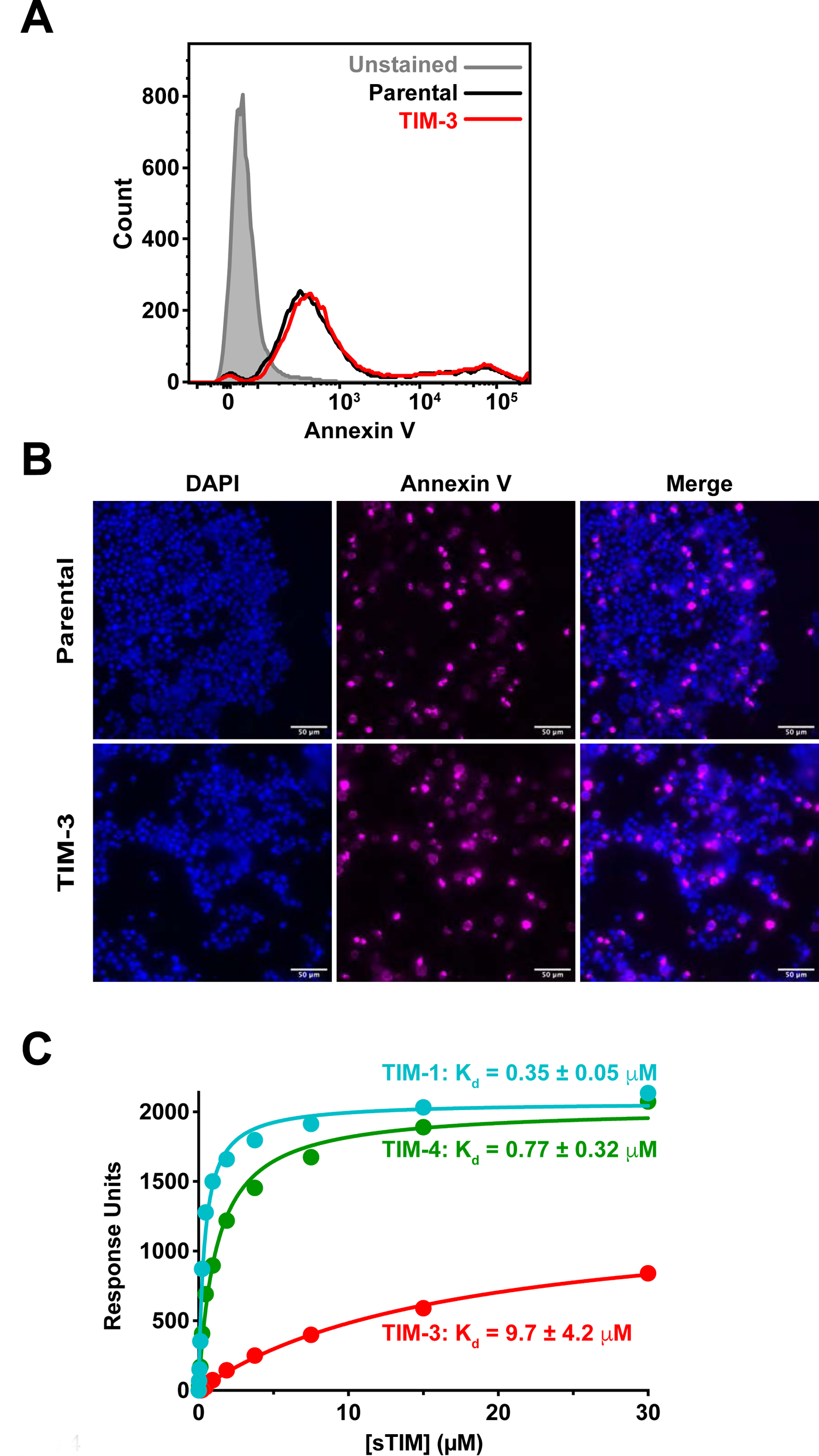
The TIM family ligand PS is present on Jurkat cells and binds TIM-3 with low micromolar affinity. (A) PS exposure on Jurkat cells in culture, detected by staining with fluorescently-tagged (PerCP-Cy^TM^5.5) Annexin V, and quantified by flow cytometry analysis of parental NF-κB GFP reporter cells (black) and those expressing TIM-3 (red) – compared with unstained cells (grey shaded). A representative histogram from 5 biological repeats is shown. (B) Fluorescence imaging of parental and TIM-3-expressing NF-κB GFP reporter cells stained with DAPI (left panels) and fluorescently-tagged (AlexaFluor 647) Annexin V (middle panels). Cells were stained with Annexin V prior to fixation and staining with DAPI. (C) PS binding of sTIM-1 (teal), sTIM-3 (red), and sTIM-4 (green) was analyzed using SPR by flowing purified TIM ECRs over 20% DOPS/80% DOPC lipid vesicles immobilized on an L1 sensorchip. Binding of TIM ECRs was determined in the presence of 1 mM CaCl_2_. The curves indicate the fit of the data in Prism 8 to a one-site specific binding equation. Binding curves are representative of at least 5 independent experiments. Mean K_d_ values and standard deviations are listed in the figure (n = 6 for sTIM-1, n = 11 for sTIM-3, and n = 5 for sTIM-4). Binding to additional lipids and effects of calcium are shown in Figure S5.

Annexin V did prove useful for assessing PS exposure on our Jurkat cells, however, and revealed that PS is indeed exposed on their surface in our cultures – presumably as cells undergo apoptosis and/or activation, two processes known to lead to PS exposure on the cell surface [27, 56, 57]. We stained cells with fluorescently labeled (PerCP-Cy^TM^5.5) annexin V, and flow cytometry analysis showed that both parental and virally-transduced TIM-3^+^ NF-κB reporter Jurkat cells expressed PS on their surface at clearly detectable levels (Figure 4A). Interestingly, most cells showed only low levels of PS exposure (peak signal ∼400), but a small population had high levels of PS exposure (∼10^5^). As a secondary approach, we also imaged cells stained with fluorescently-labeled (AlexaFluor® 647) Annexin V prior to fixation, and demonstrated Annexin V staining of both parental and TIM-3^+^ NF-κB reporter Jurkat cells (Figure 4B). These data suggest that sufficient PS may be exposed in our cellular experiments to engage TIM-3 and promote its signaling in Jurkat cells. The fact that adding PS-containing vesicles had no further influence on NF-κB activity suggests that the PS levels in culture may be near saturation and/or may represent an optimal lipid composition that is not recapitulated by our synthetic vesicles.

These data prompted us to ask whether (and how) PS functions as a TIM-3 ligand to modulate NF-κB signaling in our experiments.

### Phospholipid binding by human TIM extracellular regions

Before attempting to modulate PS binding by TIM-3 in our cellular experiments, we wanted to understand its PS-binding properties and compare them with those of TIM-1 and TIM-4, the original PS receptor [30]. We expressed the complete soluble TIM protein extracellular regions (sTIM proteins), containing the IgV and mucin domains, in Expi293 cells and used surface plasmon resonance (SPR) to measure their binding to membranes containing 20% (mole/mole) dioleoylphosphatidylserine (DOPS) in a background of dioleoylphosphatidylcholine (DOPC) as described [42]. sTIM-3 bound to these membranes with an apparent dissociation constant (K_d, app_) of 9.70 ± 4.20 μM in the presence of 1 mM CaCl_2_ (Figure 4C). The human sTIM-1 and sTIM-4 proteins bound significantly more strongly to the same membranes under identical conditions, with K_d,app_ values of 0.35 ± 0.05 μM and 0.77 ± 0.32 μM respectively (Figure 4C). The relative affinities that we observed here reflect what has been reported for murine TIM proteins using ELISA-type assays [31, 32] and other approaches [33], with sTIM-3 binding PS membranes 12-27 fold more weakly than sTIM-1 or sTIM-4. The actual K_d, app_ values that we report are substantially weaker than those suggested using ELISA methods (which were in the nM range), which is typical for multivalent systems when comparing results from direct binding studies such as SPR with those from ELISA assays [43, 58]. Our K_d, app_ values are in the same range as those measured for other PS binding proteins, pleckstrin homology domains and other peripheral membrane proteins [42, 59, 60], and indeed, for recognition of the prototypical T cell co-receptors CD28 and PD-1 for their respective ligands, CD80/86 [61] and PD-L1 [62].

In additional binding studies (Supplementary Figure 5), we further found that the presence of Ca^2+^ ions increased the PS-binding affinities of sTIM-1 and sTIM-4 by ∼10-fold, but increased the affinity of sTIM-3 for PS by just ∼2-fold (Supplementary Figure 5A, Supplementary Table 1) – consistent with previous work [33]. We also asked whether the sTIM proteins can bind other phospholipids known to become exposed on the cell surface during cell death, namely phosphatidic acid (PA) and phosphatidylethanolamine (PE) [63, 64]. sTIM-1, sTIM-3, and sTIM-4 all bound to 20% PA-containing membranes, but saturated at a slightly lower level than on membranes containing the same level of PS (Supplementary Figures 5C-E). Interestingly, inclusion of 5% PA also enhanced binding of all sTIM proteins to membranes containing 20% PS (Supplementary Figures 5F-H), such that sTIM-3 bound more strongly to membranes containing 20% PS plus 5% PA than to membranes containing 20% or 25% PS (Supplementary Figure 5I). Only sTIM-1 showed significant binding to membranes containing the zwitterionic phospholipid PE (Supplementary Figure 5C). This is consistent with a previous report [65] that also indicated that PE on the surface of apoptotic cells promotes their TIM-1-mediated phagocytosis.

**FIGURE 5.**
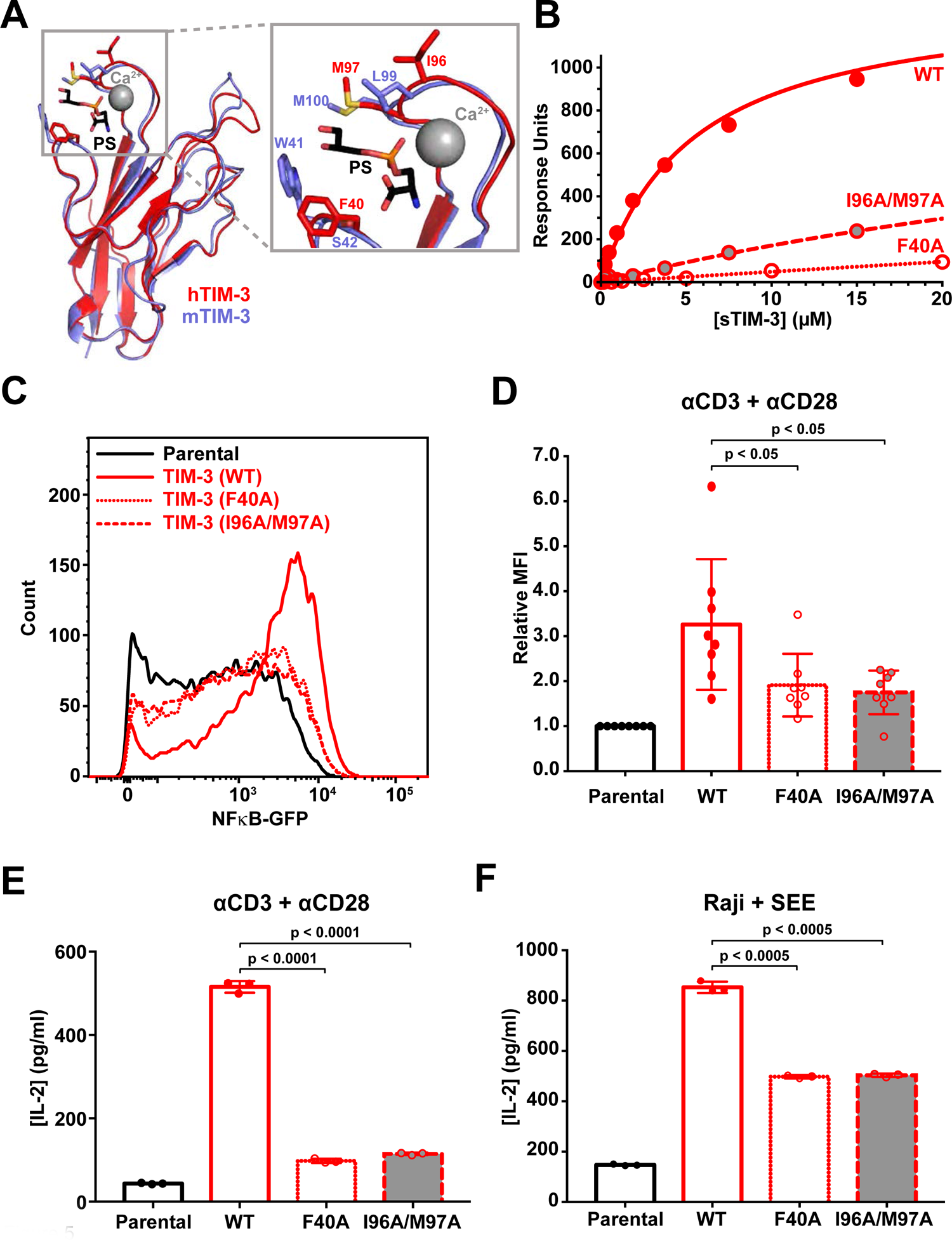
Mutated TIM-3 variants with impaired PS binding fail to activate signaling. (A) Details of the PS binding site in murine TIM-3 (PDB: 3KAA – blue) overlaid on human TIM-3 (PDB: 6TXZ – red), revealing key interacting residues [31, 80]. Inset at right shows the PS-binding pocket, where short-chain PS is sandwiched by L99/M100 and W41 in mTIM-3 (I96/M97 and F40 in human TIM-3). A calcium ion (grey sphere) also sits in the binding pocket and interacts with the negatively charged PS headgroup. (B) PS binding of wild-type sTIM-3 (red solid curve), sTIM-3^F40A^ (red dotted curve), and sTIM-3^I96A/M97A^ (red dashed curve) were analyzed using SPR by flowing the purified sTIM-3 variants over 20% DOPS/80% DOPC lipid vesicles immobilized on an L1 chip (in the presence of 1 mM CaCl_2_). Binding curves are representative of at least 3 independent experiments. K_d_ values for sTIM-3^F40A^ and sTIM-3^I96A/M97A^ were too high to measure accurately, both appearing to exceed approximately 200 μM. (C) A GFP reporter was used to measure NF-κB transcriptional activity downstream of TCR activation with 1 μg/ml αCD3 plus 0.5 μg/ml αCD28 for 16 h. A representative histogram of NF-κB-driven GFP expression is shown for parental NF-κB GFP reporter cells (black), and those expressing TIM-3^WT^ (red), TIM-3^F40A^ (red dotted), and TIM-3^I96A/M97A^ (red dashed) variants. (D) The relative mean GFP fluorescence intensity (MFI), expressed as fold change from parental cells within each experiment, was determined across 8 biological replicates for parental NF-κB GFP reporter cells (black) or those expressing TIM^WT^ (solid red line, open bar), TIM^F40A^ (dotted red line, open bar), or TIM^I96A/M97A^ (dashed red line, grey filled bar). Means ± SD are plotted, with p values determined by two-tailed, unpaired Student’s t-test. (E) ELISA analysis of IL-2 secreted into cell culture medium by parental NF-κB GFP reporter cells, and those expressing, TIM-3^WT^ (solid red line, open bar), TIM-3^F40A^ (dotted red line, open bar), or TIM-3^I96A/M97A^ (dashed red line, grey filled bar) following stimulation by αCD3/αCD28, as described in (**C**). Means ± SD are plotted for 3 biological repeats. p values comparing TIM-3^WT^ with the mutated TIM-3 variants were determined using two-tailed, unpaired Student’s t-test. (F) SEE-loaded Raji B cells were used to stimulate NF-κB GFP reporter Jurkat cells to recapitulate formation of an immunological synapse. IL-2 production following Raji B cell-based activation was determined by IL-2 ELISA analysis for parental NF-κB GFP reporter cells (solid black line, open bar) and cells expressing TIM-3^WT^ (solid red line, open bar), TIM-3^F40A^ (dotted red line, open bar), or TIM-3^I96A/M97A^ (dashed red line, grey filled bar). Mean ± SD are plotted for 3 biological repeats, with p values comparing TIM-3^WT^ with the mutated TIM-3 variants determined using two-tailed, unpaired Student’s t-test.

### Mutations that impair TIM-3 binding to PS reduce its impact on T cell signaling

Reasoning that the effect of TIM-3 expression on TCR signaling in Jurkat cells reflects TIM-3 engagement by PS on the cell surface in our experiments, we next mutated TIM-3 to reduce its PS-binding affinity. Our goal was to ask whether such mutations diminish the effect of TIM-3 on TCR signaling. Guided by the crystal structure of the IgV-like domain in the murine TIM-3 extracellular region complexed with a short acyl-chain PS [31], we substituted F40 of TIM-3 (mature human protein numbering) with alanine in one variant (F40A), and replaced both I96 and M97 with alanine in another (I96A/M97A). The corresponding residues in murine TIM-3 (W41/S42, and L99/M100 in PDB entry 3KAA) ‘sandwich’ the bound PS (Fig. 5A). DeKruyff et al. (2010) showed previously that mutating L99 and M100 of murine TIM-3 greatly impaired PS binding. They also showed that mutating W41 in mTIM-3 abrogated binding, and the W41 side-chain occupies a very similar location to that of the F40 side-chain in hTIM-3 (Figure 5A).

We first used SPR to confirm that these mutations reduce PS binding. Both sTIM-3^F40A^ and sTIM-3^I96A/M97A^ displayed substantially reduced PS binding (Figure 5B), reaching only 8% and ∼30% of saturation respectively at the highest protein concentrations tested. We generated NF-κB reporter Jurkat cells stably-expressing full length TIM-3 variants with these mutations. Having used Western blotting and flow cytometry to confirm that the mutated variants are robustly expressed (Supplementary Figure 2E) and at the cell surface (Supplementary Figure 2F), we assessed NF-κB signaling and IL-2 secretion after αCD3/αCD28 stimulation of the TCR. Cells expressing the mutated TIM-3 variants showed significantly reduced TCR-induced NF-κB activation compared with that seen for cells expressing wild-type TIM-3 (Figures 5C,D). Moreover, cells expressing the mutated variants secreted significantly less IL-2 into their medium compared with cells expressing wild-type TIM-3 (Figure 5E). Similar results were observed when using SEE-loaded Raji B cells to activate the Jurkat cells, where cells expressing mutated variants of TIM-3 produced less IL-2 than wild-type TIM-3-expressing cells (Figure 5F). These results demonstrate that reducing the ability of TIM-3 to bind PS also diminishes its ability to promote TCR signaling, supporting the hypothesis that TIM-3 is engaged with (or saturated by) PS in our Jurkat cell experiments. This in turn argues that PS can promote co-stimulatory TIM-3 signaling in this T cell model.

### A TIM-3 antibody that blocks PS binding also diminishes T cell signaling

One concern with such mutational studies is that the mutations might impair TIM-3 function indirectly, through misfolding, for example. We therefore investigated the consequences of blocking PS binding with a TIM-3 antibody. Several TIM-3 antibodies are now in clinical trials in lung cancer and other malignancies [23]. TIM-3 antibodies that have shown efficacy in preclinical studies in mice were all shown to block PS and CEACAM1 (but not galectin-9) binding [34]. The same study also showed that the human TIM-3 monoclonal antibody F38.2E2 blocks PS binding, by interacting with the loops that contain F40 and I96/M97 shown in Figure 5A, and which we mutated for the studies described above. As an orthogonal approach to testing the importance of PS binding for TIM-3 signaling, we explored the effect of F38.2E2 on co-stimulatory TIM-3 signaling, assessing both NF-κB activation and IL-2 secretion following TCR stimulation in our Jurkat reporter cell assay. As CEACAM1 is not endogenously expressed in Jurkat cells [66, 67], any observed effects from F38.2E2 treatment should be specific to blocking the ability of TIM-3 to bind PS.

F38.2E2 addition caused a modest, but significant, dose-dependent decrease in αCD3/αCD28-stimulated NF-κB signaling in cells expressing TIM-3, but not in parental cells (Figures 6A-E). The effect of F38.2E2 treatment became saturated at approximately 1 μg/ml antibody (Figures 6B,E). Importantly, 10 μg/ml F38.2E2 also reduced (by ∼50%) the amount of IL-2 secreted by TIM-3 expressing cells upon TCR activation (Figure 6F). Since F38.2E2 is known to block PS binding, these data provide additional support for the argument that TIM-3 engagement by PS is important for its co-stimulatory signaling activity in this system. Inhibition is only partial, but F38.2E2 blocks ∼50% of the TIM-3-dependent *increase* in NF-κB signaling or IL-2 secretion, bringing the relative MFI from ∼1.0 to <0.7 for TIM-3 cells compared with a value of ∼0.35 for parental cells (compare Figures 6D and E), and reducing IL-2 levels from ∼500 to ∼300 pg/ml (compared with ∼40 pg/ml for parental cells). One possible explanation for the fact that the observed inhibition is partial is that ligands other than PS are also important – such as galectin-9, binding of which to TIM-3 is not blocked by F38.2E2. Alternatively, the partial nature of the response could reflect the need for F38.2E2 to compete with a high level of PS on cells in culture, which might be clustered as suggested on activated T cells [56] in a way that enhances its avidity for TIM-3 binding. F38.2E2 antibody treatment itself could also cross-link or cluster TIM-3 in a way that modulates the avidity of TIM-3 for PS, reducing its inhibitory effect. In either case, taken together with the mutational and chimeric receptor studies, our data strongly support a role for extracellular PS binding in regulating TIM-3 co-stimulatory signaling when overexpressed in Jurkat cells, and suggest that inhibition of this signaling is an important component of the function of TIM-3 antibodies currently being studied in the clinic.

**FIGURE 6.**
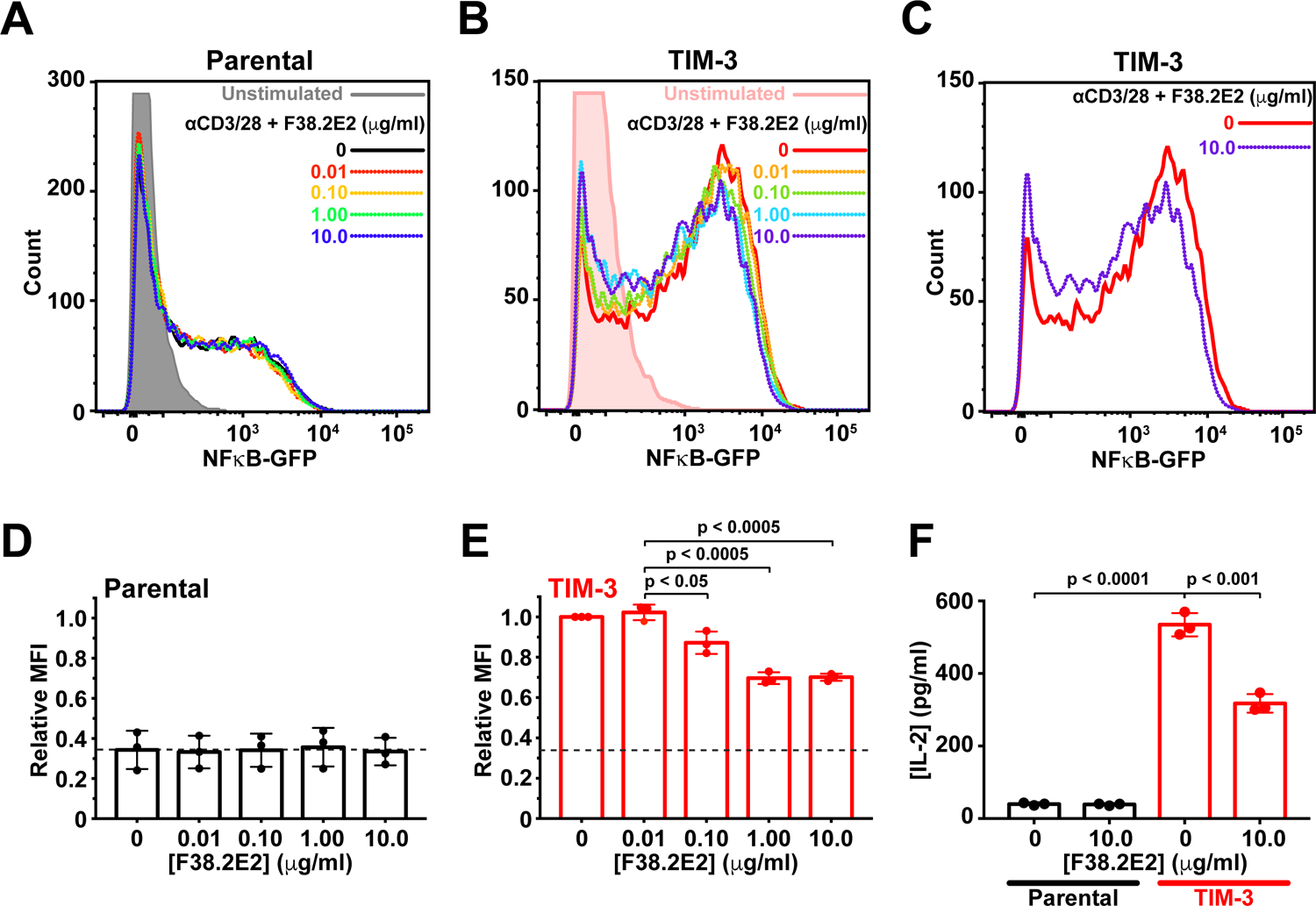
TIM-3 antibody treatment selectively reduces TIM-3 signaling. (A) Representative histograms of NF-κB-driven GFP expression in parental NF-κB GFP reporter Jurkat cells were starved for 4 h – including a 1 h pre-treatment with anti-TIM3 (F38.2E2) at the concentrations marked – and then stimulated with 1 μg/ml αCD3 plus 0.5 μg/ml αCD28 for 16 h or left unstimulated (grey). GFP expression was analyzed by flow cytometry, and quantitation is shown in (**D**). (B) Representative histograms for TIM-3-expressing NF-κB GFP reporter Jurkat cells, stimulated as in (**A**), with 0 μg/ml to 10 μg/ml F38.2E2, and unstimulated TIM-3^+^ cells shown in pink. Quantitation is shown in (**E**). (C) Representative histograms from TIM-3-expressiong NF-κB GFP reporter Jurkat cells stimulated with 0 μg/ml (red) or 10 μg/ml F38.2E2 (blue), showing that 10 μg/ml F38.2E2 does reduce the level of TCR-induced NF-κB-driven GFP expression seen in stimulated TIM-3 cells. (**D**-**E**) Quantitation of data in (**A**) and (**C**) over 3 independent biological repeats, plotting mean values for relative mean GFP fluorescence intensity (MFI) and standard deviations. p values were determined with two-tailed, unpaired Student’s t-tests. (**F**) IL-2 secreted into cell medium was measured by ELISA for parental NF-κB GFP reporter Jurkat cells (black bars) and TIM-3-expressing cells (red bars). Adding 10 μg/ml F38.2E2 reduces enhanced IL-2 production in TIM-3 expressing cells. Bars represent mean concentration of IL-2 (± SD) for 3 biological replicates. p values were determined with two-tailed, unpaired Student’s t-tests.

## DISCUSSION

Our findings provide evidence that PS binding to the extracellular region of TIM-3 directly modulates TIM-3 co-signaling activity in a Jurkat T cell model. Consistent with several previous studies, TIM-3 expression in Jurkat cells had a co-stimulatory effect on TCR signaling. We find that this effect requires the intact extracellular region of TIM-3 or another (PS-binding) TIM family member. Moreover, the co-stimulatory effect of TIM-3 is abolished by mutations that impair PS binding – or by treatment with an antibody that occludes the PS-binding site on TIM-3. These results argue that PS exposed on the surface of Jurkat cells in culture – which we confirm with annexin V staining – is sufficient to engage TIM-3 and promote its co-stimulatory signaling. The role of PS binding in regulating signaling effects of TIM-3 has remained largely unexplored [8, 68, 69]. PS binding to TIM-3 on macrophages, dendritic cells and fibroblasts has been shown to mediate their phagocytosis of apoptotic cells [31, 32, 70], whereas the same interaction only allows T cells to form conjugates with apoptotic cells – with no engulfment [31]. Our studies argue that, beyond this coupling function, PS binding to TIM-3 can regulate its activity as a TCR co-signaling receptor.

Several previous studies have reported the presence of a TIM-3 ligand on the surface of various cultured cells, from CD4 T cells [13, 14] to fibroblasts to macrophages – and from several different species [55]. The cross-species reactivity observed supports the suggestion that this ligand is PS rather than a protein, and Cao et al. (2007) demonstrated that it is not galectin-9, the most well-studied potential TIM-3 ligand. Together with our results and existing evidence for PS binding by TIM-3 [31–33], it seems likely that most published studies of TIM-3 signaling in culture have been performed with TIM-3 already engaged by a ligand (PS). Nonetheless, the prevailing hypothesis in the field is that galectin-9 is the key TIM-3 ligand and binds TIM-3 to trigger co-inhibitory signaling and thus T cell death [8, 24, 71]. However, as a carbohydrate-binding protein, galectin-9 is known to bind numerous other targets [72–75]. Moreover, several studies have argued that some effects of galectin-9 on T cells are TIM-3-independent [26, 75, 76] or alternatively that galectin-9 does not bind TIM-3 at all [25]. One interesting possibility is that PS is in fact the primary ligand for TIM-3, and that galectin-9 modulates PS binding and/or its consequences.

Alternatively, our findings may suggest that the inhibitory signaling reported for TIM-3 reflects displacement of an activating ligand – namely PS. Galectin-9 could exert its TIM-3-dependent effects by pulling TIM-3 into clusters that no longer bind PS, or bind it in a different way. Lee et al. (2011), commented in studies of TIM-3 co-stimulatory signaling that the receptor initially augments T cell activation, but that rapid inhibition of T cell activation is seen when TIM-3 is crosslinked with an antibody. Interestingly, that antibody was 5D12, which has subsequently been shown to block PS binding to TIM-3 [34] – so this phenomenon could actually have arisen through reversal of PS stimulation. TIM-3 has also been reported to enhance DC and NK cell activation, as well as FcεR1 signaling in mast cells [15, 77, 78]. Moreover, stimulation of TIM-3 with PS was recently reported to promote its phosphorylation in natural killer (NK) cells, leading to inhibition of cytokine secretion [79]. It therefore seems clear that TIM-3 can signal in several different contexts, possibly exerting distinct effects (and effects in different directions) through differential ligand engagement and/or phosphorylation of the tyrosines in its cytoplasmic tail.

Several groups are currently pursuing development of therapeutic antibodies against TIM-3 as a novel immune checkpoint strategy, mostly in combination with PD-1 or PD-L1 antibodies [3, 8, 23]. Identification of the biologically relevant ligand(s) is essential for identifying candidate therapies most likely to succeed in the clinic. The findings described here support the importance of PS as a key signaling ligand for TIM-3. Crucially, the epitopes of most functional TIM-3 antibodies have been shown to map to the PS binding site [34, 80] but do not affect galectin-9 binding, arguing that blocking PS binding is key to conferring functional efficacy of the antibodies in promoting immune responses. Most TIM-3 antibodies currently in the clinic block PS binding to the receptor [3], including Sym023 [81], BGB-A425 [82], ICAGN02390 [83], and LY3321367 [84]. Our findings argue that these antibodies most likely block a signaling response to PS binding. Understanding TIM-3 signaling that elicits this response, and the context(s) in which it can be beneficial, will be important for furthering our understanding of this unusual TCR co-signaling receptor, and for understanding potential side-effects of TIM-3 blockade – which have already been seen in some preclinical contexts [85].

## DATA AVAILABILITY

The data needed to evaluate this work are included in the manuscript, and are available upon request.

## COMPETEING INTERESTS

The authors declare no competing interests related to this study.

## AUTHOR CONTRIBUTIONS

C.M.S. and M.A.L. designed the overall project and wrote the manuscript. C.M.S. performed all experiments and wrote the first manuscript draft. A.L. and N.K. assisted in experiments with TIM-3 chimerae. All authors contributed to analysis of results and manuscript editing.

## ACKNOWLEDGEMENTS

We thank members of the Lemmon and Ferguson laboratories, as well as Aaron Ring and Carla Rothlin for valuable discussions and comments.

