## Supplementary Figures 1-5 and Table S1 for "Phosphatidylserine binding regulates TIM-3 effects on T cell receptor signaling"

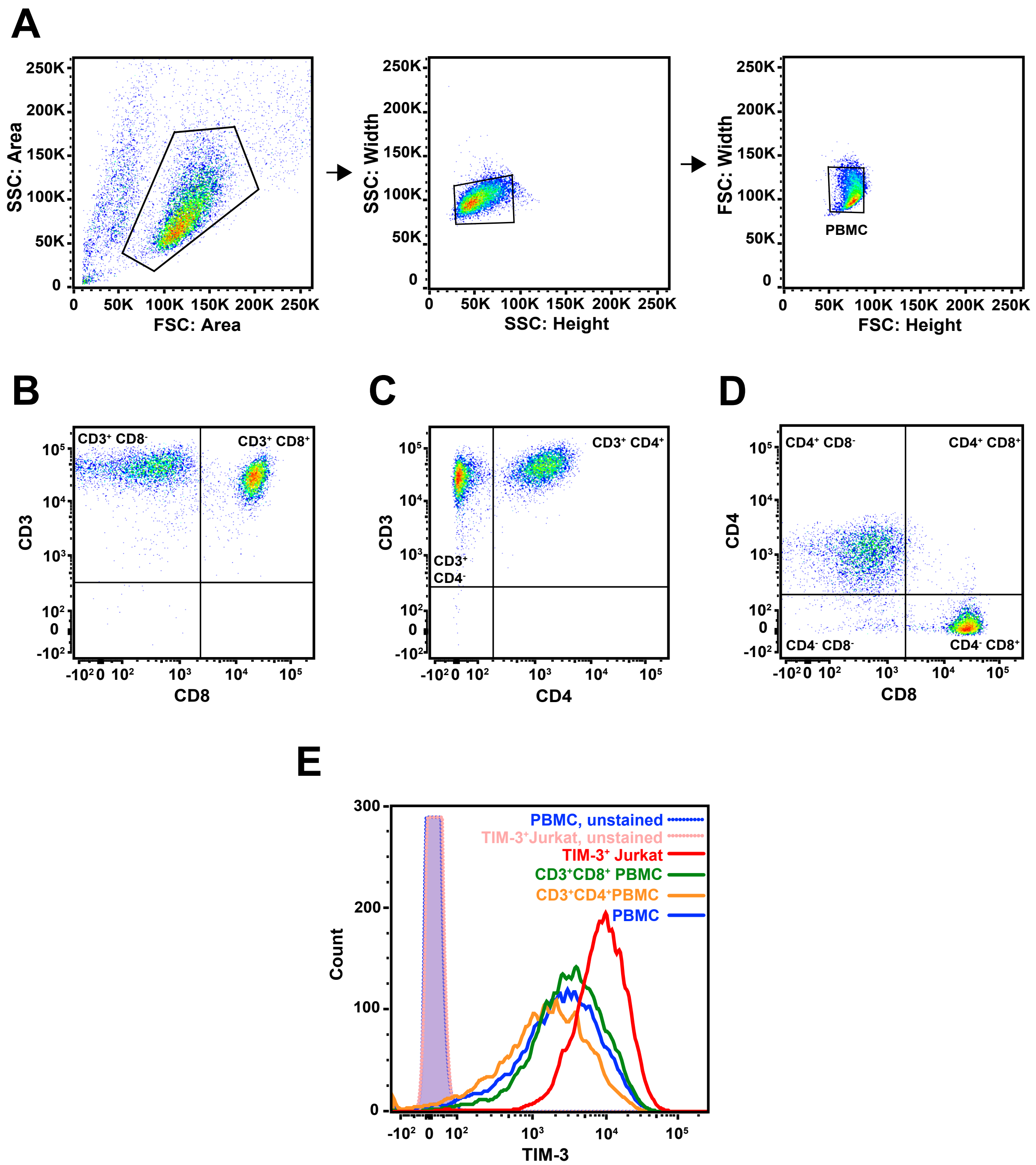


**Supplementary Figure 1.**
**Comparison of TIM-3 expression in primary human T cells and virally-transduced Jurkat NF-κB reporter cells**

After 7 days of stimulation with αCD3/αCD28, human peripheral blood mononuclear cells (PBMCs) were analyzed by flow cytometry for expression of CD3, CD8, CD4, and TIM-3.

(**A**) Doublet discrimination was performed to select singlet PBMCs for analysis. Detected events were analyzed and gated as shown (left panel). Gated cells were assessed for their side scatter height and width (middle panel), and then forward scatter height and width (right panel) to discriminate doublets and select singlets, in gate termed "PBMC".

(**B-D**) Analysis of CD3, CD8, and CD4 expression in the PBMC population confirm that T cells were expanded during 7-day stimulation, as the majority of cells express CD3 (**B, C**). Approximately half of the $CD3^+$ cells express CD8 or CD4, as shown in the $CD3^+CD8^+$ or $CD3^+CD4^+$ in panels (**B**) and (**C**), respectively. (**D**) The expression of CD8 and CD4 is mutually exclusive, as expected.

**(E**) Expression of TIM-3 in TIM-3-expressing Jurkat NF-κB cells (red) is compared to TIM-3 expressed by $CD3^+CD8^+$ PBMCs (green), $CD3^+CD4^+$ PBMCs (orange), or the bulk PBMC population (blue), as defined in dot plots in (**B**), (**C**), and (**A**, right panel), respectively. Unstained PBMC (blue) and $TIM\text{-}3^+$ Jurkat NF-κB cells (pink) controls are shown in dotted lines with shaded peaks.

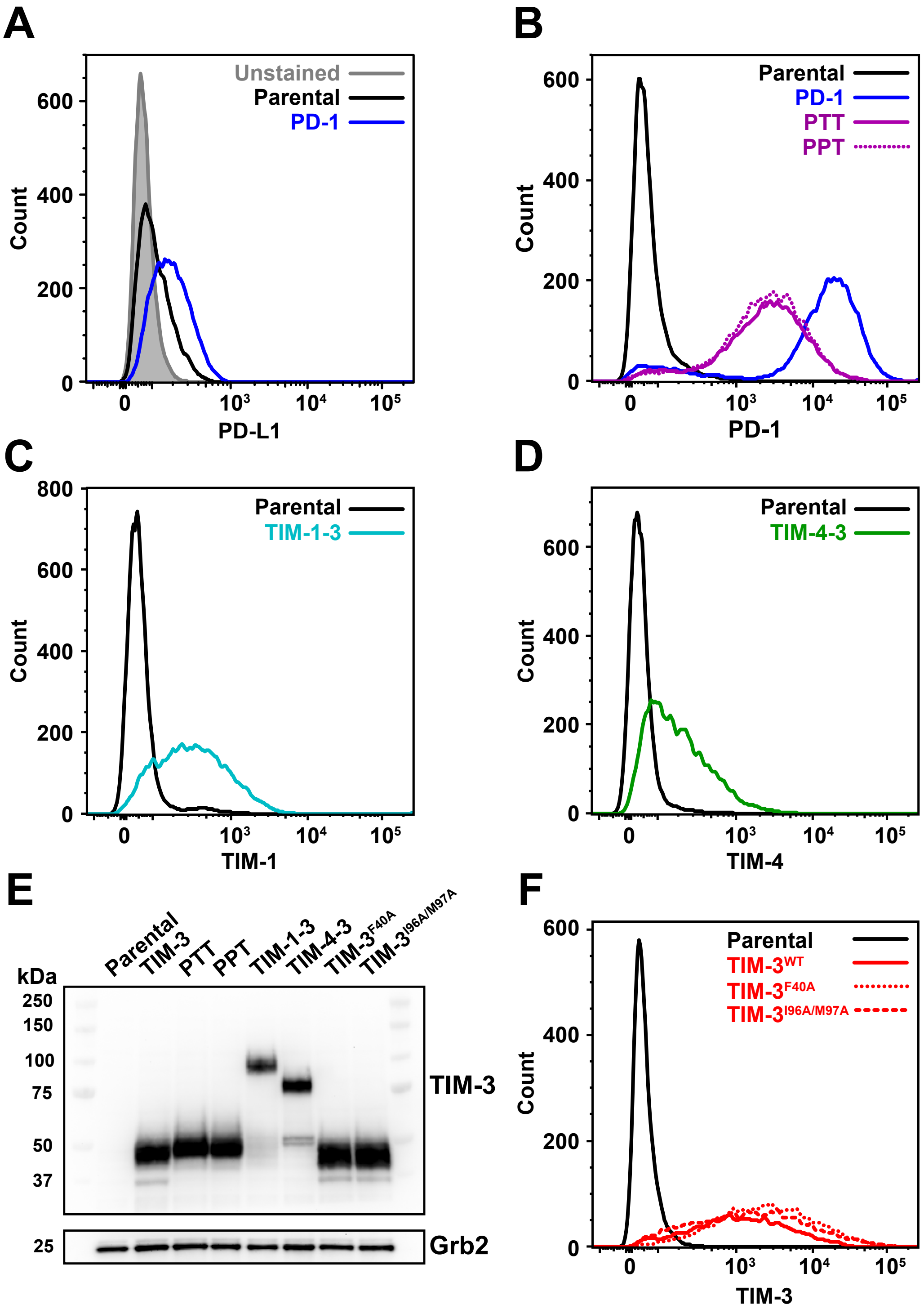


**Supplementary Figure 2.**
**Expression of receptors in Jurkat cells after lentiviral transduction**

(**A**) NF-κB GFP transcriptional reporter Jurkat cells were analyzed for PD-L1 expression by flow cytometry. Parental (black curve) or PD-1-expressing (blue curve) cells were stained with fluorescently labeled anti-PD-L1 and compared to unstained PD-1-expressing cells (grey shaded).

(**B**) Parental NF-κB reporter Jurkat cells do not express detectable levels of PD-1. Lentivirus transduction was used to exogenously express PD-1 or the PD-1/TIM-3 chimerae (PTT and PPT) in NF-κB reporter Jurkat cells. Surface receptor expression was confirmed by staining with fluorescently labeled anti-PD-1, with representative histograms shown.

(**C**, **D**) Jurkat cells do not express TIM-1 (**C**) or TIM-4 (**D**), but do express the chimeric TIM-1-3 and TIM-4-3 chimerae when introduced by lentiviral transduction (**C**, **D**).

(**E**) Chimeric receptors and mutated TIM-3 variant expression in lentivirus-transduced NF-κB reporter Jurkat cells are similar to full-length, wild-type TIM-3, assessed by Western blotting with an antibody against the intracellular region of TIM-3.

(**F**) Mutated TIM-3 variants with defective PS binding (TIM-$3^{F40A}$ and TIM-$3^{I96A/M97A}$) were robustly expressed at the cell surface following lentiviral transduction, as detected by staining with anti-TIM-3 and labeled anti-goat antibody to ensure staining of all TIM-3 variants.

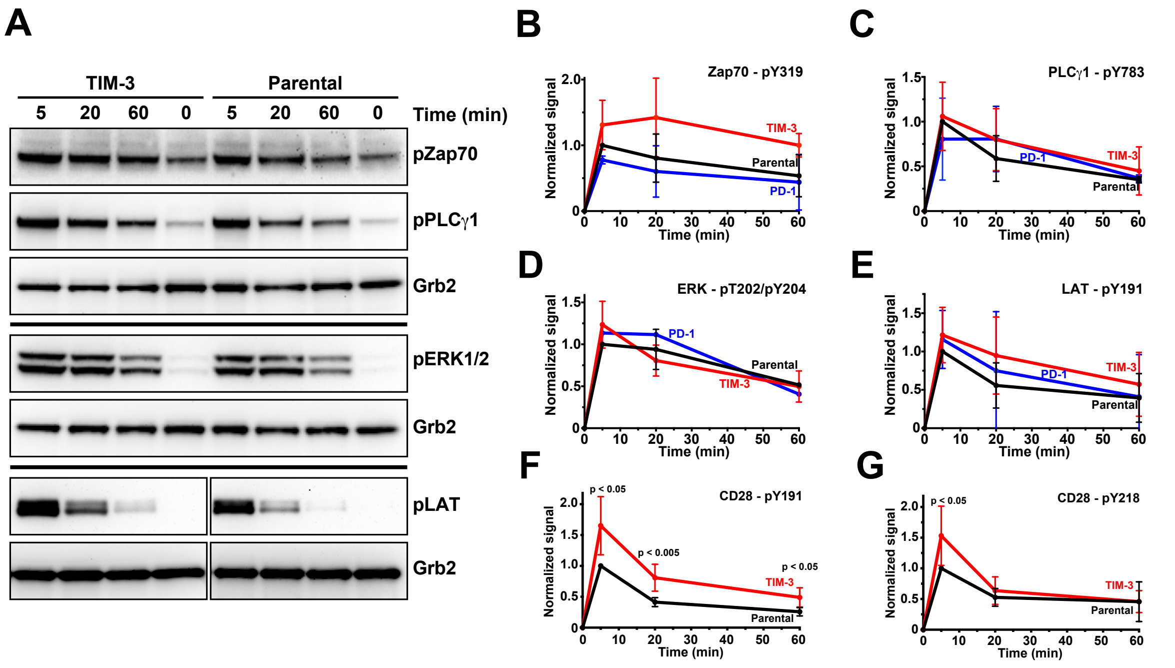


**Supplementary Figure 3.**
**Effects of TIM-3 expression on TCR signaling components**

Western blotting was used to monitor changes in phosphorylation of TCR-associated signaling molecules following TCR activation. Parental NF-κB GFP reporter Jurkat cells (black), or those expressing TIM-3 (red) or PD-1 (blue) were starved for 4 h, and then stimulated with 1 μg/ml αCD3 plus 1 μg/ml αCD28 for 0, 5, 20, or 60 minutes

(**A**) Representative Western blot with phospho-specific antibodies for Zap70 (pY319), PLCγ1 (pY783), ERK1/2 (pT202/pY204), and LAT (pY191), with Grb2 as a loading control, for TIM-3-expressing cells (left lanes) and parental NF-κB reporter Jurkat cells (right lanes).

(**B**-**E**) Band intensities for (**B**) phospho-Zap70 (pY319), (**C**) phospho-PLCγ1 (pY783), (**D**) phospho-ERK1/2 (pT202/pY204), and (**E**) phospho-LAT (pY191) are shown after 5, 20, and 60 min of stimulation, quantified using the Kodak ImageStation and normalized to Grb2. Data points for Zap70, LAT, and PLCγ1 represent mean values for at least 3 independent experiments (2 for PD-1). Data points for pERK1/2 represent mean values for 2 independent experiments for parental and TIM-3 cells and 1 experiment for PD-1. Error bars show standard deviation. Statistical analysis with two-tailed, unpaired Student's t-test to compare values for TIM-3 and PD-1 cells to parental cells showed no significant differences at any time point.

(**F**-**G**) Quantification of phosphoCD28 at (**F**) Y191 and (**G**) Y218, with representative blots shown in **Figure 2A,B**, for 5, 20, and 60 min of stimulation. Bands were quantified using the Kodak ImageStation and normalized to Grb2. Points represent the average of 5 repeats for each phosphosite with standard deviation shown in error bars. p values comparing TIM-3 and parental cells were determined with a two-tailed, unpaired Student's t-test.

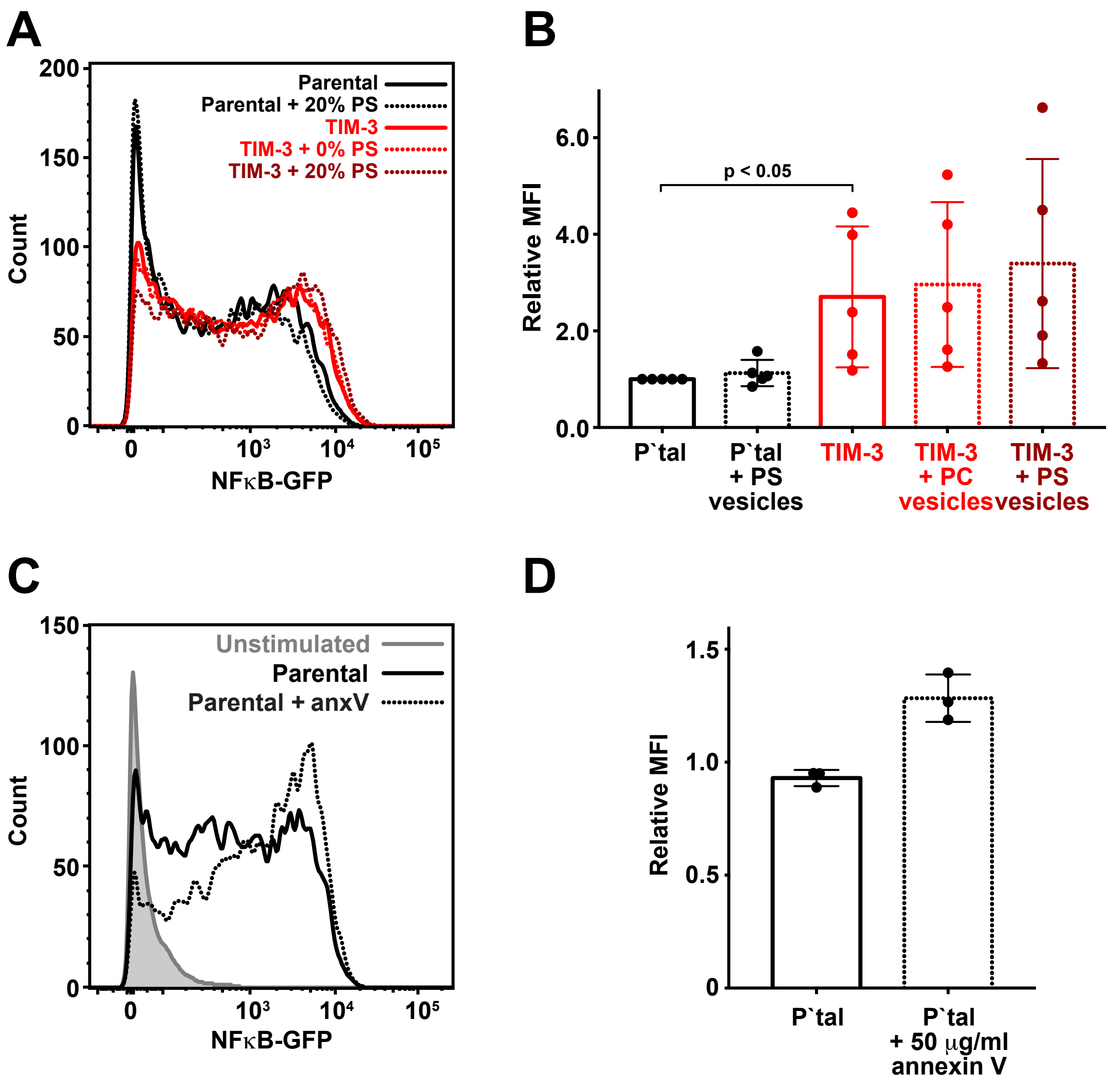


**Supplementary Figure 4.**
**Addition of exogenous PS fails to increase NF-κB-driven transcription in TIM-3⁺ cells**

(**A**) Representative histograms of NF-κB-driven GFP expression in parental NF-κB GFP reporter Jurkat cells (solid black curve) that had been serum starved for 4 h, and then stimulated with 1 μg/ml αCD3 plus 0.5 μg/ml αCD28 for 16 h. Equivalent experiments were performed in parallel with these parental cells treated with 100 μM 20% DOPS/80% DOPC vesicles (black dotted curve), TIM-3-expressing cells (red solid curve), TIM-3-expressing cells treated with 100 μM 100% DOPC vesicles (red dotted curve), and TIM-3-expressing cells treated with 100 μM 20% DOPS/80%DOPC vesicles (dark red dotted curve). Vesicle addition to TIM-3-expressing cells has no significant influence, as quantitated across experiments in (**B**).

(**B**) Quantitation of relative mean GFP fluorescence intensity (MFI) from repeats of the experiments in (**A**), showing substantial spread in the data, but no significant difference between treatments across 5 independent repeats. Bars represent mean ± SD.

(**C**) Representative histograms of NF-κB-driven GFP expression in unstimulated (grey, shaded) and stimulated parental NF-κB GFP reporter cells alone (black, solid line) or with 50 μg/ml Annexin V (anxV) (black, dotted line), showing that Annexin V promotes GFP expression in this system even in stimulated parental cells.

(**D**) Quantitation of data in (**C**), comparing parental (P'tal) cells with- and without Annexin V. Mean ± SD is shown for biological triplicate.

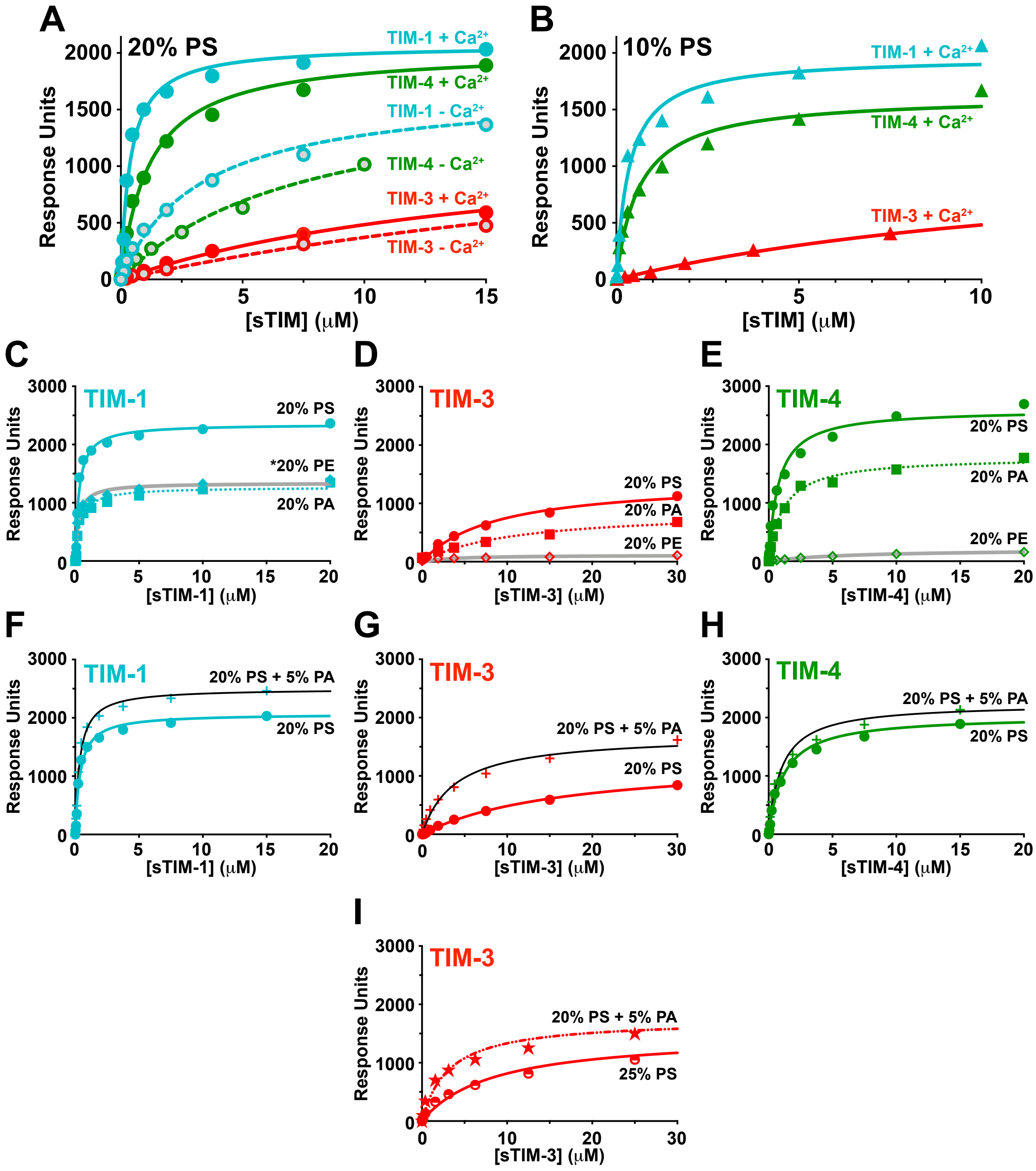


**Supplementary Figure 5.**
**Phospholipid-binding characteristics of human TIMs**

(**A**) Comparison of PS binding by sTIM-1 (teal), sTIM-3 (red), and sTIM-4 (green) with- and without 1 mM $CaCl_2$, assessed using SPR by flowing purified TIM extracellular regions (ECRs) over lipid vesicles containing 20% DOPS/80% DOPC immobilized on an L1 chip. The curves represent fits of the data to a one-site specific binding equation in Prism 8. Mean $K_d$ values and standard deviations are shown in Supplemental Table 1.

(**B**) sTIM protein binding to 10% DOPS/90% DOPC surface membranes was determined in the presence of 1 mM $CaCl_2$. Binding curves in this experiment are representative of at least 2 independent experiments, except for TIM-1 (n=1).

(**C**-**E**) sTIM binding to 20% DOPS/80% DOPC (circles), 20% DOPA/80% DOPC (squares), or 20% DOPE/80% DOPC (diamonds) surface was determined in the presence of 1 mM $CaCl_2$ for sTIM-1 (teal), sTIM-3 (red), and sTIM-4 (green), with binding curves representative of at least 2 independent experiments, except for TIM-4. Note that all sTIMs bind PA significantly, but that sTIM-1 is unique in also binding substantially to PE as described in the main text (grey line in **C**).

(**F**-**H**) sTIM binding to 20% DOPS/80% DOPC (circles) or 20% DOPS/5%DOPA/75% DOPC (crosses) surface was determined in the presence of 1 mM $CaCl_2$. For each sTIM protein, adding 5% PA (mole/mole) detectably increases binding.

(**I**) sTIM-3 binding to 20% DOPS/5% PA/75% DOPC (stars) surface was determined in the presence of 1 mM $CaCl_2$, and found to exceed that seen for 25% DOPS/75% DOPC (half-filled circles), indicating some preference for PA.

**Supplementary Table 1. SPR Data for TIM family members binding to PS-containing membranes.**

| % PS | Sample | $CaCl_2$ (mM) | $K_{d, app}$ ± Std. Dev (µM) | N |
|---|---|---|---|---|
| 10% | sTIM-1 | 1 | 0.49 | 1 |
| | sTIM-3 | 1 | 22.4 ± 12.6 | 6 |
| | sTIM-4 | 1 | 0.69 ± 0.05 | 2 |
| 20 % | sTIM-1 | 0 | 3.13 ± 0.31 | 2 |
| | sTIM-3 | 0 | 20.4 ± 10.5 | 3 |
| | sTIM-4 | 0 | 7.21 ± 0.45 | 2 |
| 20% | sTIM-1 | 1 | 0.35 ± 0.05 | 6 |
| | sTIM-3 | 1 | 9.70 ± 4.20 | 11 |
| | sTIM-4 | 1 | 0.77 ± 0.32 | 5 |
| 20 % + 5 % PA | sTIM-1 | 1 | 0.35 | 1 |
| | sTIM-3 | 1 | 3.45 ± 0.49 | 2 |
| | sTIM-4 | 1 | 1.00 | 1 |
